# Oral delivery of mesenchymal stem cell-derived extracellular vesicles to treat intestinal inflammation

**DOI:** 10.1101/2025.10.31.685898

**Authors:** Mona Belaid, Wei Heng Chng, Ram Pravin Kumar Muthuramalingam, Yun Wei Lim, Jana Javorovic, Yunyue Zhang, Xiang Luo, Bertrand Czarny, Driton Vllasaliu

## Abstract

Despite advances in therapy for inflammatory bowel disease (IBD), current treatments are still associated with poor clinical outcomes and severe systemic side effects. Extracellular vesicles derived from mesenchymal stem cells (MSC-EVs) could have therapeutic applications in IBD due to their regenerative potential and immunomodulatory properties. Previous studies investigating the potential of MSC-EVs in IBD have largely administered the vesicles by injection, which does not offer the significant benefits of oral administration, including direct and localised access to the site(s) of intestinal inflammation. Here, we evaluated the stability of MSC-EVs for oral delivery by assessing particle size, concentration and EV markers. The EVs disintegrated in gastrointestinal (GI) fluids, with cryogenic electron microscopy confirming the loss of structural integrity. To address this, we developed a double-coating formulation consisting of chitosan and Eudragit S100 to enhance GI stability and facilitate colon-targeted delivery. We demonstrated that coated EVs were resistant to GI fluids and digestive enzymes, and the formulation released intact vesicles in colonic fluid. Preliminary *in vivo* studies in a dextran sodium sulfate-induced colitis mouse model showed that coated EVs reduced disease severity, whereas uncoated EVs performed similarly to the control group. Interestingly, coated MSC-EVs elicited a stronger therapeutic response compared to EVs administered intravenously at the same dose. These findings indicate that oral delivery of MSC-EVs could be an effective route of administration, with appropriate formulation, to treat intestinal inflammation.

## Introduction

Inflammatory bowel disease (IBD) is a chronic and incurable inflammatory condition of the gastrointestinal tract. Damage to the intestinal mucosa due to infectious agents, toxins, environmental factors or an imbalance in the regulating components of the mucosal barrier can result in dysfunction and loss of barrier integrity [1, 2]. This disruption affects the host-microbial balance, leading to dysbiosis, and activates the immune system, triggering inflammatory signalling cascades that result in intestinal inflammation [3, 4]. The two major classifications of IBD include Crohn’s disease and ulcerative colitis. IBD emerged as a global public health challenge at the turn of the 21st century [5, 6], with an estimated 7 million people affected worldwide [7, 8], and its prevalence is expected to continue rising over the next two decades [9, 10]. IBD poses a substantial burden on global healthcare systems and society, resulting in high medical expenses, lost productivity and a reduced quality of life for affected individuals [5, 11, 12]. While biological therapies have improved the management of IBD, they are administered systemically by injection and are associated with high cost, toxicity and loss of therapeutic response over time [13–15]. Despite the availability of advanced treatment options, many individuals with IBD continue to experience suboptimal disease control and substantial gaps remain in the appropriate management of the condition [16]. There is therefore an unmet need for novel, safe and effective IBD therapies.

Extracellular vesicles (EVs) are cell-derived membrane-bound nanoparticles that mediate intercellular communication, elicit functional responses and promote phenotypic changes that influence the cells’ physiological or pathological status [17]. Cells release EVs containing membrane and cytosolic components with specific lipid, protein and nucleic acid compositions that reflect their biogenesis and determine their biological role [18–21]. Extracellular vesicles derived from mesenchymal stem cells (MSC-EVs) have been investigated for their therapeutic potential in IBD owing to their inherent regenerative and immunomodulatory properties. In IBD mouse models, MSC-EVs have been shown to promote mucosal healing and regeneration of the intestinal epithelium, restore barrier integrity, suppress pro-inflammatory cytokine production, reduce macrophage infiltration in colon tissues and promote M2 macrophage polarisation, and improve gut microbiota composition [22–31].

However, current studies with MSC-EVs in IBD animal models mainly use intravenous or intraperitoneal injections for the administration of EVs *in vivo* [32]. Systemically administered EVs are rapidly cleared from the bloodstream, primarily by the mononuclear phagocyte system, limiting their bioavailability and therapeutic efficacy at the site of action [33]. As a result, systemic administration of EVs often requires higher doses, which can potentially increase the risk of off-target effects as well as treatment costs. Effective treatment of IBD requires targeted delivery of therapeutics to the intestine [34] and the oral route is considered both the most suitable for achieving localised intestinal delivery and the most convenient for patients [35]. Oral administration of MSC-EVs would overcome the issues associated with intravenous administration and eliminate the requirement for a trained healthcare professional to administer the therapy.

In this study, we investigated the stability of MSC-EVs in gastrointestinal fluids and developed a coating strategy to enhance their oral stability and enable colon-targeted delivery. We then conducted preliminary *in vivo* studies to evaluate the therapeutic efficacy of orally administered coated and uncoated MSC-EVs, as well as intravenously administered EVs, in a mouse model of colitis.

## Results and Discussion

### Biophysical and biochemical characterisation of MSC-EVs

EVs were isolated from the conditioned medium of human bone marrow mesenchymal stem cells. The size distribution and concentration of EVs were measured using nanoparticle tracking analysis (Figure 1a). 90% of isolated EVs were smaller than 200 nm in diameter and most batches had a particle to protein ratio (P:P) > 3×10^10^ particles/µg, which indicated high vesicular purity [36] (Figure 1b). The EVs had a mean surface charge of −17 mV as determined by zeta potential measurements. Images captured by cryogenic electron microscopy (cryo-EM) showed the EV morphology as spherical membrane-bound vesicles with surface structures (Figure 1c). EV-specific markers were detected using the Exo-Check^TM^ exosome antibody array kit and the vesicles showed a positive expression for transmembrane proteins CD63 and CD81 and cytosolic proteins TSG101, ALIX and FLOT1, and a negative expression for cis-Golgi matrix protein GM130, which confirmed the absence of cellular contamination in the isolated EV preparation [37, 38] (Figure 1d). Super-resolution microscopy also confirmed the colocalization of tetraspanins CD63, CD81 and CD9 on single EVs (Figure 1e).

**Figure 1.**
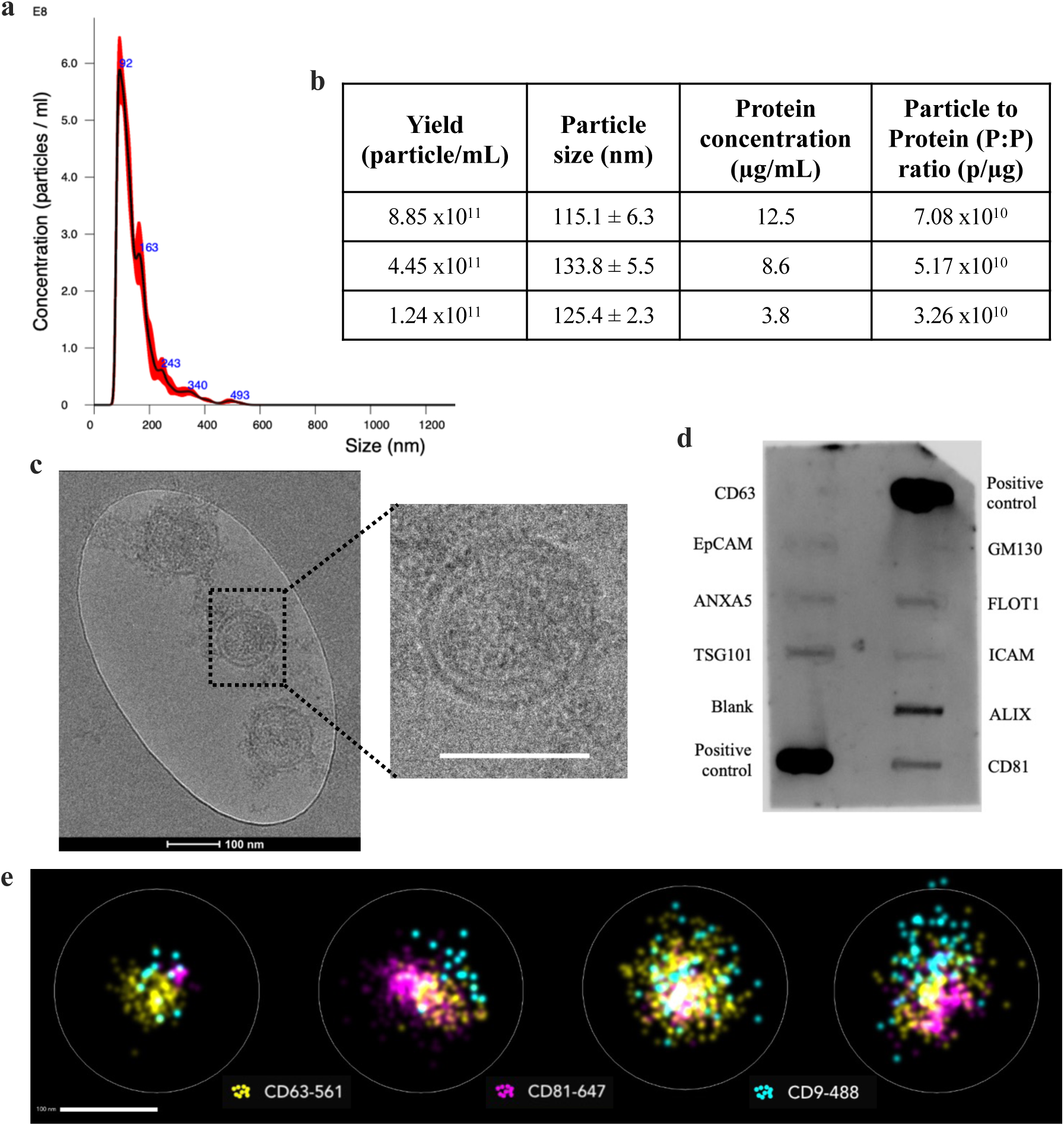
Biophysical and biochemical characterisation of MSC-EVs. **(a)** Size distribution of MSC-EVs using nanoparticle tracking analysis (NTA). The NTA histogram represents different captures with red areas denoting standard deviation between measurements. **(b)** Table summarising the characterisation of different EV preparations. Yield and particle size were measured by NTA. Protein concentration was measured using QuantiPro^TM^ bicinchoninic acid (BCA) assay. The particle to protein (P:P) ratio was calculated using the formula: P:P ratio = Yield / Protein concentration. **(c)** Cryogenic electron microscopy (cryo-EM) images of MSC-EVs showing spherical vesicular morphology with a bilayer membrane. Scale bar: 100 nm. **(d)** Detection of EV markers using Exo-Check^TM^ exosome antibody array kit. CD63 and CD81: tetraspanins, EpCAM: epithelial cell adhesion molecule, ANXA5: annexin A5, TSG101: tumour susceptibility gene 101, FLOT1: flotillin-1, ICAM: intercellular adhesion molecule 1 and ALIX: programmed cell death 6 interacting protein. The membrane also includes four controls, GM130: cis-Golgi matrix protein as a negative marker for cellular contamination during EV isolation, two positive controls and one blank. **(e)** Super-resolution microscopy images of MSC-EVs stained with three tetraspanins: CD63 (yellow), CD81 (pink) and CD9 (blue). Scale bar: 100 nm.

### MSC-EVs possess regenerative and immunomodulatory properties

To assess cellular responses to MSC-EVs, we performed *in vitro* assays using three concentrations: 10^7^, 10^9^ and 10^11^ particles per mL (p/mL).

We examined the effects of MSC-EVs on metabolic activity and proliferation in Caco-2 epithelial cells. Cells incubated with EVs at 10^7^ and 10^9^ p/mL for 4 days showed a significant increase in cell density, whereas 10^11^ p/mL EVs produced results similar to the control group (Figure 2a). None of the EV concentrations diminished Caco-2 cell viability, unlike the strong cytotoxic effect observed with 10% dimethyl sulfoxide (DMSO). Caco-2 cells seeded in culture inserts with a cell-free gap and incubated with MSC-EVs were imaged every 24 h to monitor wound healing (Figure 2b). Epithelial wound repair requires a combination of cell proliferation and migration [39]. EVs at 10^9^ p/mL significantly increased the percentage of wound closure at 48 h and 72 h (Figure 2c) and the rate of cell migration at 24 h (Figure 2d), outperforming the other EV concentrations. EVs at 10^7^ p/mL also significantly increased wound closure percentage at 72 h but did not affect the rate of Caco-2 cell migration, while 10^11^ p/mL EVs were comparable to the control group.

**Figure 2.**
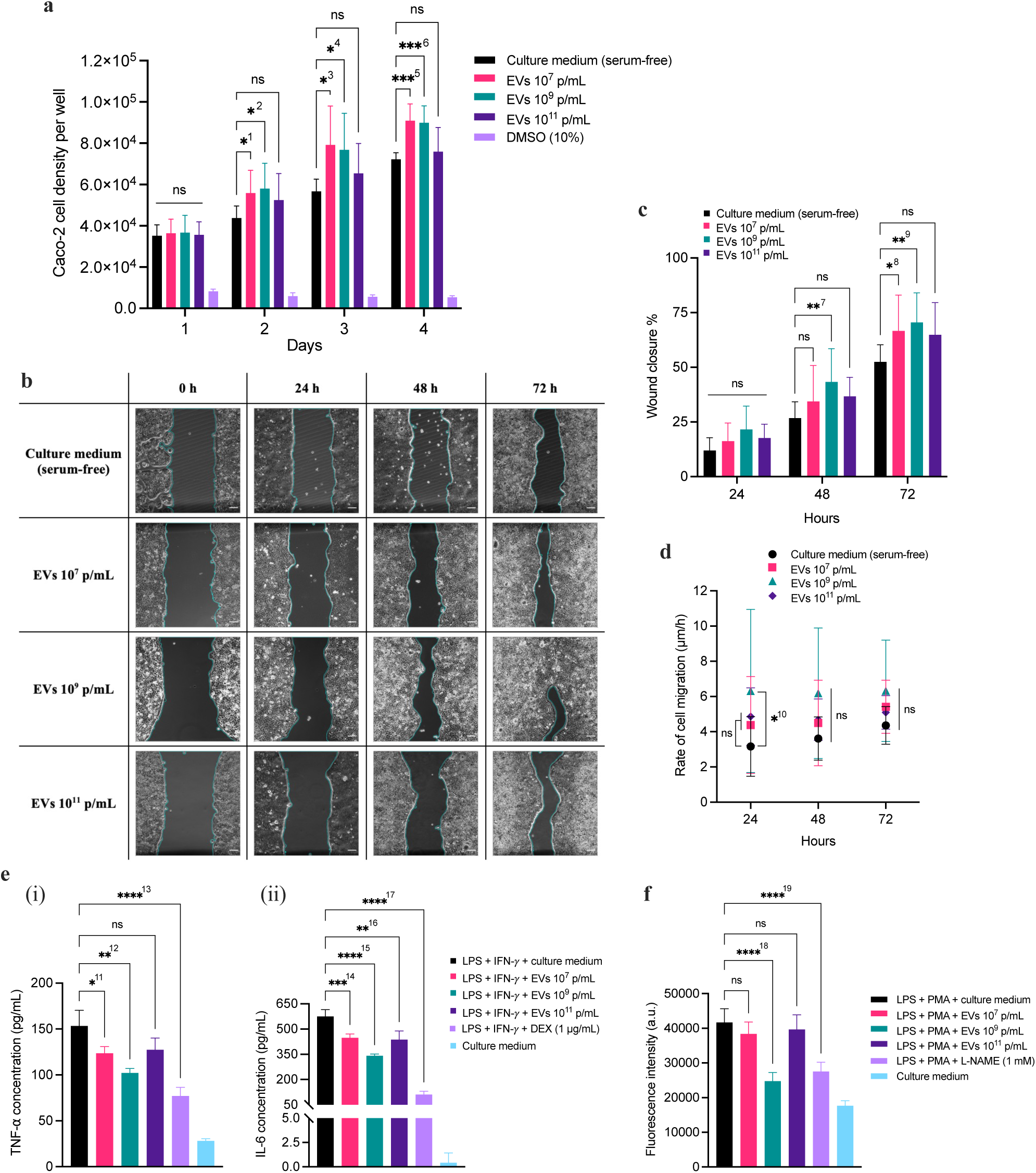
Regenerative and immunomodulatory properties of MSC-EVs. Assays were performed with three EV concentrations: 10^7^, 10^9^ and 10^11^ p/mL **(a)** Effect of MSC-EVs on cell proliferation. Caco-2 cell density per well was obtained using the MTS assay standard curve (Supplementary Figure 2). EVs at 10^7^ and 10^9^ p/mL significantly increased Caco-2 cell density per well on day 2 (^1^* p = 0.0391, ^2^* p = 0.0287), day 3 (^3^* p = 0.0219, ^4^* p = 0.0296) and day 4 (^5^*** p = 0.0002, ^6^*** p = 0.0004). 10% Dimethyl sulfoxide (DMSO) was used as a negative control for cell proliferation. **(b)** Wound healing images of Caco-2 cells captured every 24 h and analysed using Image J to measure wound area and width. Scale bar: 100 µm. **(c)** Effect of MSC-EVs on wound closure percentage over time. 10^9^ p/mL EVs significantly accelerated Caco-2 wound closure after 48 h (^7^* p = 0.0092) and 72 h (^9^** p = 0.0040). 10^7^ p/mL EVs showed a significant increase in wound closure after 72 h (^8^* p = 0.0309). **(d)** Effect of MSC-EVs on the rate of cell migration. 10^9^ p/mL EVs significantly increased the rate of Caco-2 cell migration after 24 h (^10^* p = 0.0181). **(e)** Effect of MSC-EVs on cytokine production: **(i)** TNF-⍺ and **(ii)** IL-6. J774A.1 macrophages were activated with lipopolysaccharide (LPS) and IFN-𝛾 and cytokine levels were quantified using the Luminex assay. EVs at 10^7^ and 10^9^ p/mL significantly decreased the production of TNF-α (^11^* p = 0.0242, ^12^** p = 0.0017) and IL-6 (^14^*** p = 0.0008, ^15^**** p < 0.0001). EVs 10^11^ p/mL did not affect TNF-α but did reduce the concentration of IL-6 (^16^** p = 0.0029). 1 µg/mL Dexamethasone (DEX) was used as a positive control for anti-inflammatory activity and significantly inhibited the production of both TNF-α and IL-6 (^13^**** and ^17^**** p < 0.0001). **(f)** Effect of MSC-EVs on oxidative stress. J774A.1 macrophages were stimulated with LPS and phorbol 12-myristate 13-acetate (PMA), and reactive oxygen species (ROS) were detected by measuring fluorescence intensity following DCFH-DA staining. 10^9^ p/mL was the only EV concentration that significantly decreased ROS production (^18^**** p < 0.0001). L-NAME: Nω-Nitro-L-arginine methyl ester. 1 mmol/L L-NAME was used as a positive control for antioxidant activity and significantly reduced oxidative stress (^19^**** p < 0.0001). Data are presented as mean ± SD (n = 3). **(a)**, **(c)** and **(d)** Two-way ANOVA with post-hoc Dunnett’s test. **(e)** and **(f)** Brown-Forsythe and Welch ANOVA with post-hoc Dunnett’s T3 test. (* p < 0.05, ** p < 0.01, *** p < 0.001, **** p < 0.0001).

We evaluated the effects of MSC-EVs on cytokine production and oxidative stress during inflammation in J774A.1 macrophages. Lipopolysaccharide (LPS) stimulation in the presence of IFN-𝛾 polarises macrophages toward the M1 phenotype, which is characterised by the production of high levels of pro-inflammatory cytokines, such as TNF-α and IL-6 [40]. Activated J774A.1 cells incubated with EVs at 10^7^ p/mL and 10^9^ p/mL showed a significant reduction in TNF-α levels after 24 h (Figure 2e (i)). All three EV concentrations significantly lowered IL-6 levels after 24 h, with 10^9^ p/mL EVs exerting the strongest effect (Figure 2e (ii)). Exposure to LPS and phorbol 12-myristate 13-acetate (PMA) induces cellular oxidative stress, as pathogen-derived compounds increase the production of reactive oxygen and nitrogen species (ROS and RNS, respectively) in macrophages [41]. Only 10^9^ p/mL EVs significantly reduced ROS production in J774A.1 macrophages after 24 h, whereas 10^7^ and 10^11^ p/mL EVs produced similar outcomes to the control group (Figure 2f).

These results demonstrated that the most effective EV dose *in vitro* was 10^9^ p/mL. At this concentration, MSC-EVs enhanced metabolic activity, proliferation and wound healing in epithelial cells, together with reducing cytokine production and oxidative stress in inflamed macrophages. While EVs at 10^7^ p/mL elicited positive effects on cell proliferation, wound healing and pro-inflammatory cytokine release, they did not affect ROS production and were overall less effective than 10^9^ p/mL EVs. By comparison, EVs at 10^11^ p/mL had little impact on cells apart from reducing IL-6 secretion, possibly because they were degraded, as high EV doses can up-regulate lysosomal activity [42].

### MSC-EVs reduce inflammation and restore epithelial barrier integrity in a co-culture model of intestinal inflammation

We investigated the effect of MSC-EVs in a Caco-2/J774A.1 co-culture model that mimics intestinal inflammation [43]. The model consists of human Caco-2 intestinal epithelial cells and murine J774A.1 macrophages cultured in a Transwell^®^ system, which comprises a semipermeable membrane insert separating the apical (upper) and basolateral (lower) compartments. Caco-2 epithelial cells were seeded on the Transwell^®^ inserts and maintained in culture for 19-21 days until they differentiated into polarised intestinal epithelial monolayers. On the same day, differentiated Caco-2 cells were primed with a cytokine cocktail (TNF-α, IFN-γ and IL-1β), and J774A.1 macrophages were seeded and stimulated with LPS and IFN-𝛾. After 24 h, the two cell lines were combined into a co-culture, with differentiated Caco-2 epithelial cells in the apical compartment and J774A.1 macrophages in the basolateral compartment. This inflamed co-culture was characterised by elevated levels of the pro-inflammatory cytokines TNF-α and IL-6, and by loss of epithelial barrier integrity, as measured by transepithelial electrical resistance (TEER).

MSC-EVs (10^9^ particles/mL) were applied to the inflamed co-culture either in the basolateral compartment, facing activated J774A.1 macrophages, or in the apical compartment, facing the primed Caco-2 monolayer (Figure 3a). Basolateral application represents intravenous administration where EVs reach the subepithelial space (including immune cells) from the circulation, whereas apical application models oral administration where EVs access the intestinal epithelium from the luminal side. Both applications induced comparable effects in the inflamed co-culture. After 6 h of incubation, MSC-EVs significantly reduced TNF-α levels 2-fold (Figure 3b) and IL-6 levels 3-fold (Figure 3c) in the basolateral compartment, and this effect persisted after 12 h. EVs applied apically crossed the Caco-2 intestinal epithelium (Supplementary Figure 5) and exerted anti-inflammatory activity equivalent to EVs applied directly to macrophages basolaterally. MSC-EVs were also internalised by differentiated Caco-2 cells (Supplementary Figure 6) and J774A.1 macrophages (Supplementary Figure 7). After 12 h of EV treatment, TEER values began to increase in the intestinal epithelial monolayers and were restored to within 10% of healthy co-culture values by 16 h (Figure 3d). These results demonstrated that apical administration of EVs reduced macrophage cytokine production and re-established epithelial barrier integrity to the same extent as basolateral application. This finding suggests that oral delivery of MSC-EVs could be an effective approach to treat intestinal inflammation.

**Figure 3.**
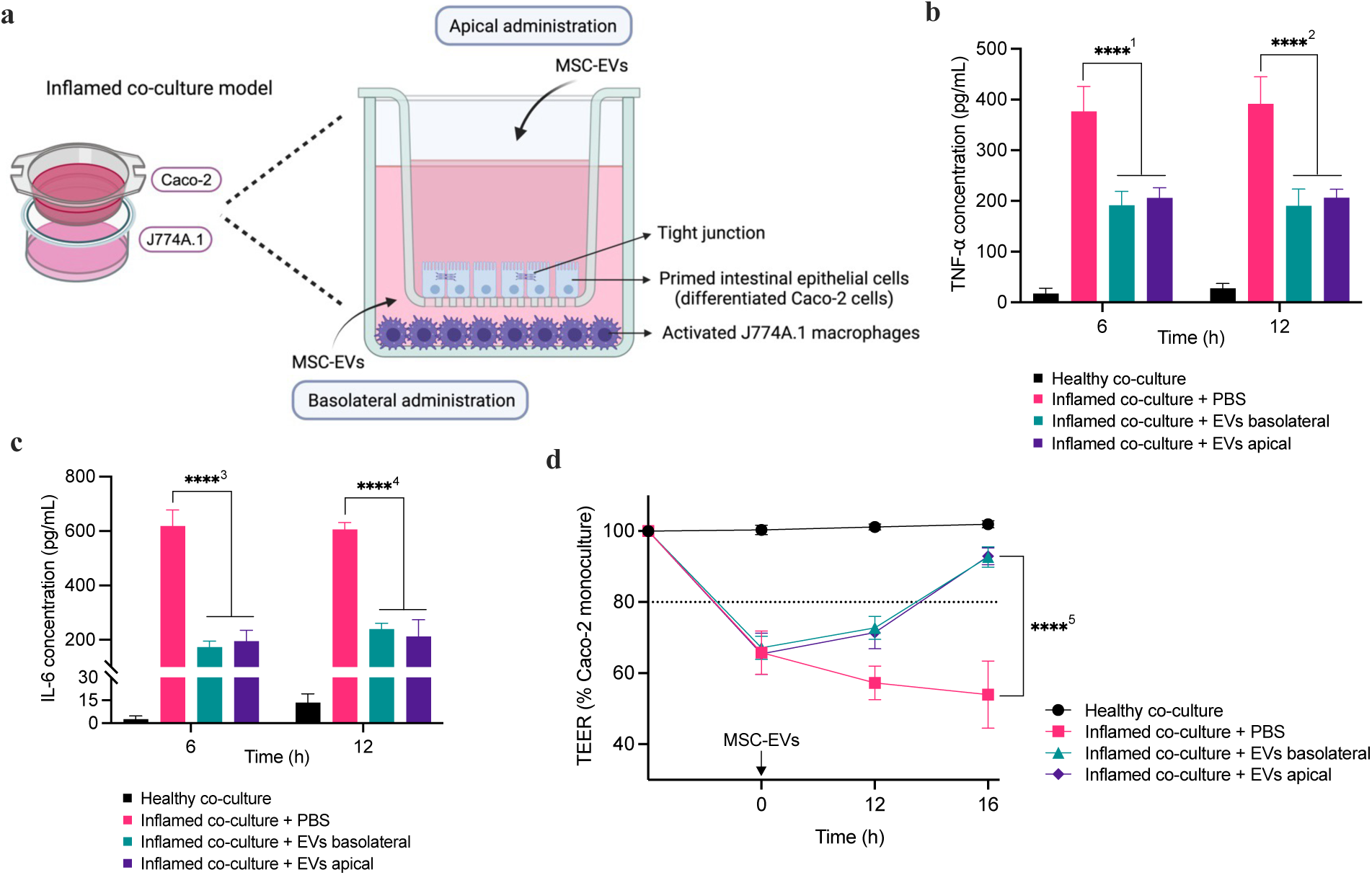
Effect of MSC-EVs in a co-culture model that mimics intestinal inflammation. **(a)** The inflamed co-culture consists of differentiated Caco-2 cells primed with cytokines (TNF-α, IFN-γ and IL-1β) and J774A.1 macrophages activated with LPS and IFN-γ in a Transwell^®^ cell culture system. MSC-EVs were applied to the co-culture either in the apical (upper) or basolateral (lower) compartment. Created with BioRender.com. Both EV applications significantly reduced the production of **(a)** TNF-α and **(b)** IL-6 in macrophages after 6 h (^1^**** and ^3^**** p < 0.0001) and 12 h (^2^**** and ^4^**** p < 0.0001). **(c)** Epithelial barrier integrity measured as transepithelial electrical resistance (TEER). A characteristic feature of the inflamed co-culture is a TEER reduction greater than 20% of the baseline Caco-2 monoculture values (0 h). MSC-EVs applied to the apical and basolateral compartments of the co-culture significantly increased TEER values after 16 h (^5^**** p < 0.0001). Data are presented as mean ± SD (n = 3). Two-way ANOVA with post-hoc Dunnett’s test. (**** p-value < 0.0001).

### MSC-EVs are not stable in gastrointestinal fluids

Stability in the gastrointestinal tract is an essential prerequisite for oral administration. To investigate the stability of MSC-EVs for oral delivery, we first incubated the EVs with simulated gastric fluid (GF), followed by simulated intestinal fluid (IF) to mimic digestion *in vitro*.

EVs exposed to GF plus pepsin and IF plus pancreatin showed a significant decrease in particle size distribution, as indicated by mean size, mode, and percentiles D10, D50, and D90 (Figure 4a). MSC-EVs incubated with GF and IF also exhibited a significant reduction in particle concentration (4-fold) (Figure 4b) and in the percentage of EV subpopulations positive for TSG101 (2.5-fold) and CD81 (8-fold) (Figure 4c). Nanoflow cytometry scatter plots showed fewer detected events and a higher proportion of unstained nanoparticles in GF and IF (94.3%) than EVs in water (61.4%) (Figure 4d). Simulated gastrointestinal fluids had a disruptive effect on EVs comparable to sodium dodecyl sulfate (SDS), a surfactant that induces vesicle lysis. MSC-EVs were also visualised at different stages of incubation with GF and IF in the presence of digestive enzymes using cryogenic electron microscopy (Figure 4e). After 30 min, structural changes were evident and vesicles appeared non-spherical with disrupted or broken bilayer membranes (Figure 4e (i) and (ii)). After 1 h, vesicles had completely lost structural integrity and images revealed distorted membrane remnants (Figure 4e (iii)). Collectively, these results demonstrate that MSC-EVs are disintegrated in gastrointestinal fluids and are therefore not stable for oral administration without appropriate formulation.

**Figure 4.**
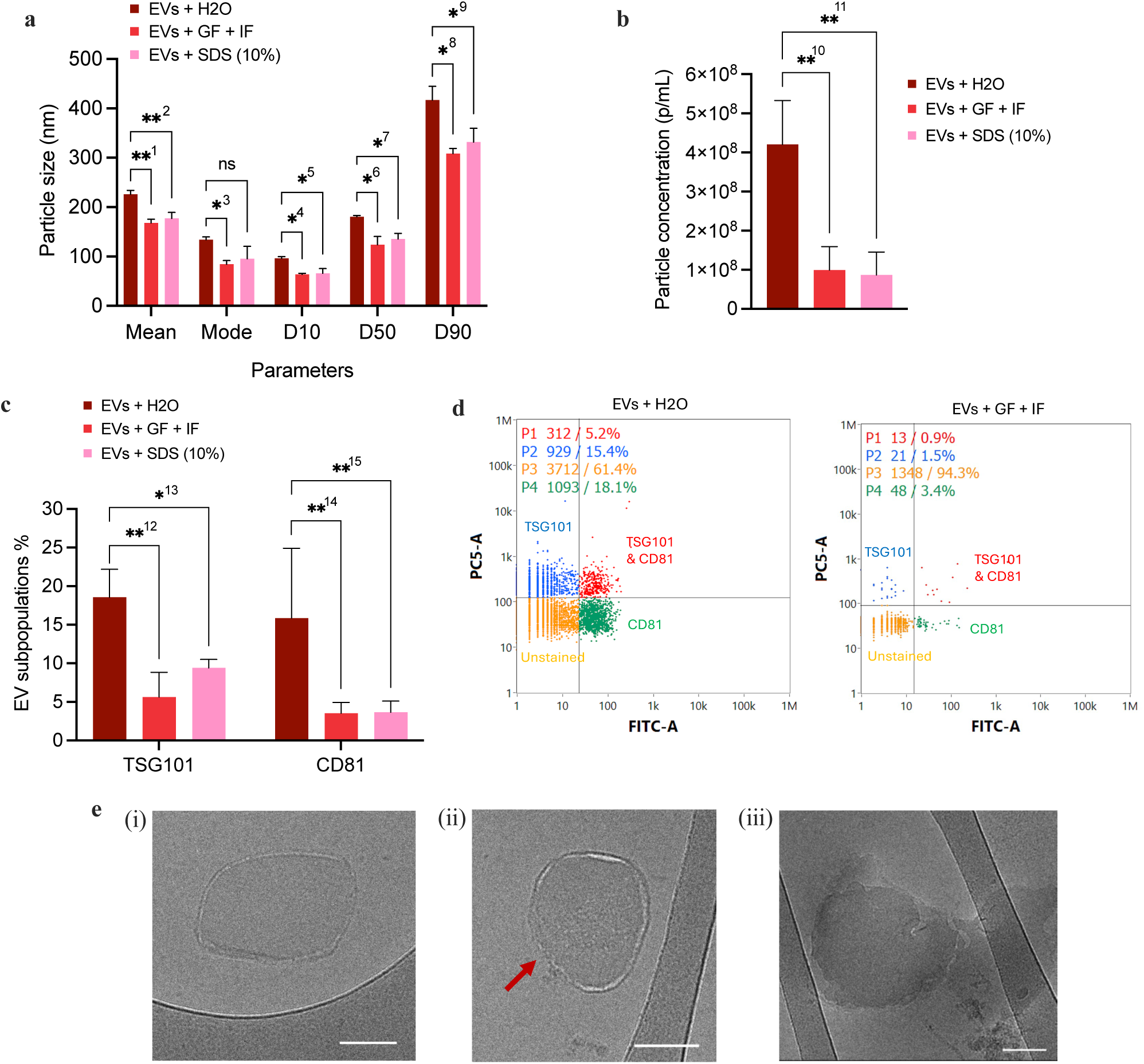
Stability of MSC-EVs in gastrointestinal fluids. MSC-EVs were incubated in simulated gastric fluid (GF) followed by simulated intestinal fluid (IF) to mimic digestion in vitro. **(a)** Particle size distribution of MSC-EVs measured by NTA. All size parameters significantly decreased after incubation with GF and pepsin + IF and pancreatin: mean (^1^** p = 0.0020), mode (^3^* p = 0.0153), D10 (^4^* p = 0.0119), D50 (^6^* p = 0.0361) and D90 (^8^* p = 0.0227). 10% sodium dodecyl sulfate (SDS) was used as a positive control for EV disintegration and showed significant reduction in mean (^2^** p = 0.0089), D10 (^5^* p = 0.0431), D50 (^7^* p = 0.0249) and D90 (^9^* p = 0.0454). **(b)** Particle concentration of MSC-EVs measured by nanoflow cytometry. Incubation with GF and IF significantly reduced the particle concentration of EVs (^10^** p = 0.0052) to the same extent as 10% SDS (^11^** p = 0.0043). **(c)** Percentage of EV subpopulations positive for TSG101 and CD81 relative to total particles detected by nanoflow cytometry. Incubation of EVs with GF and IF significantly reduced the percentage of EVs expressing cytosolic protein TSG101 (^12^** p = 0.0057) and transmembrane protein CD81 (^14^** p = 0.0078). 10% SDS also significantly reduced TSG101-positive EVs (^13^* p = 0.0407) and CD81-postive EVs (^15^** p = 0.0084). **(d)** Representative dot plots showing gated populations of stained EVs based on fluorescence intensities, FITC-A: anti-CD81 (488) vs. PC5-A: anti-TSG101 (647), measured with flow nanoanalyzer (NanoFCM). P1: Double-positive for TSG101 and CD81; P2: TSG101-only positive; P3: Double-negative (unstained); P4: CD81-only positive. The subpopulation of EVs positive for TSG101 and/or CD81 decreased following incubation with GF and IF. **(e)** Cryogenic electron microscopy (cryo-EM) images of MSC-EVs at different stages of incubation with GF and IF and digestive enzymes pepsin and pancreatin. Scale bar: 100 nm. After 30 min, **(i)** the vesicle morphology is no longer spherical and **(ii)** the bilayer membrane is ruptured (red arrow). **(iii)** After 1 h, the vesicle has completely lost its structural integrity, leaving behind irregular membrane debris. Data are presented as mean ± SD (n = 3). **(a)** and **(c)** Two-way ANOVA with post-hoc Dunnett’s test. **(b)** One-way ANOVA with post-hoc Dunnett’s test. (* p-value < 0.05, ** p-value < 0.01).

### Double-coating formulation improves oral stability of MSC-EVs and targets colon delivery

To facilitate oral delivery of MSC-EVs, we developed a double-coating formulation to protect the vesicles from degradation in the gastrointestinal (GI) environment and to target delivery to the colon. MSC-EVs were sequentially coated with chitosan as the first layer and Eudragit S-100 as the second layer to form chitosan–Eudragit-coated EVs, referred to as coated EVs (Figure 5a). Chitosan coating of EVs is driven by electrostatic attraction between the positively charged chitosan and the negatively charged EV surface. Following coating, the total number of particles in the solution was preserved, and the chitosan-coated EVs exhibited a positive surface charge and an increase in average particle size by 1.5–2-fold (Figure 5b). The structural differences between uncoated and chitosan-coated EVs were also visualised by cryogenic electron microscopy (Figure 5c (i) and (ii), respectively). When anionic Eudragit S-100 is deposited onto chitosan-coated EVs, electrostatic attraction between the oppositely charged polymers leads to the formation of a polyelectrolyte complex on the EV surface. This interaction results in the assembly of a solid double-layer coating around the vesicles, causing the coated EVs to precipitate from the solution. Scanning electron microscopy (SEM) images revealed individually coated particles embedded within a surrounding matrix (Figure 5c (iii) and Supplementary Figure 10a).

**Figure 5.**
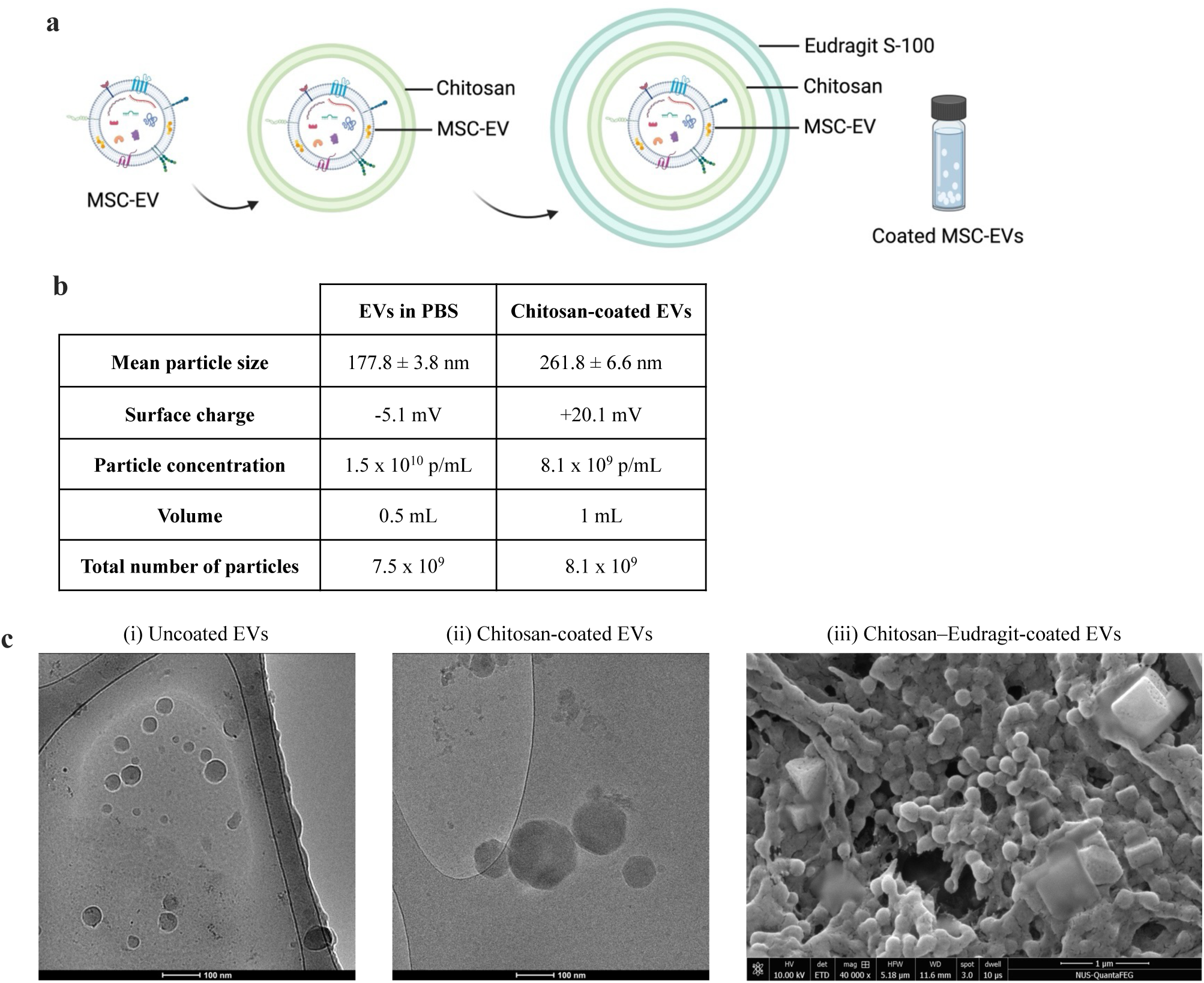
Double-coating formulation of MSC-EVs. **(a)** Schematic representation of the coating process. MSC-EVs were sequentially coated with chitosan (first layer) and Eudragit S-100 (second layer) to form chitosan–Eudragit-coated EVs, referred to as coated EVs. Cationic chitosan adheres to negatively charged EVs, then anionic Eudragit binds to the chitosan layer, thereby forming a layer-by-layer structure via electrostatic attraction. Created using BioRender.com. **(b)** Comparison between uncoated EVs and chitosan-coated EVs. Particle size and concentration were analysed using nanoparticle tracking analysis (NTA), and surface charge was measured by zeta potential. **(c)** Microscopy images showing each step of the coating process. (i) Uncoated EVs and (ii) chitosan-coated EVs were visualised by cryogenic electron microscopy (cryo-EM). Scale bar: 100 nm. (iii) Chitosan–Eudragit-coated EVs (referred to as coated EVs) were imaged with scanning electron microscopy (SEM). Scale bar: 1 µm.

The double-coating formulation employs a sequential pH- and bacteria-responsive system to enable targeted delivery to the colon. Chitosan was selected as the inner layer because it is specifically degraded by microbial enzymes produced by colonic bacteria. Its mucoadhesive properties [44] may also enhance retention and release of EVs on mucosal surfaces of the colon. As a cationic polymer, chitosan stabilises the coating by acting as an intermediate electrostatic layer between the negatively charged EV surface and the anionic Eudragit, promoting the assembly of a stable double-layer structure. Eudragit S-100 was selected as the outer layer due to its resistance to pH conditions in the stomach and proximal small intestine. The methacrylate copolymer is widely used in enteric coatings as it only dissolves at pH above 7.0, enabling drug release in the ileocolonic region [45], which can increase the local bioavailability of EVs in the colon. The stability of coated EVs in the upper GI tract and colon-targeted release of EVs are illustrated in Figure 6a.

**Figure 6.**
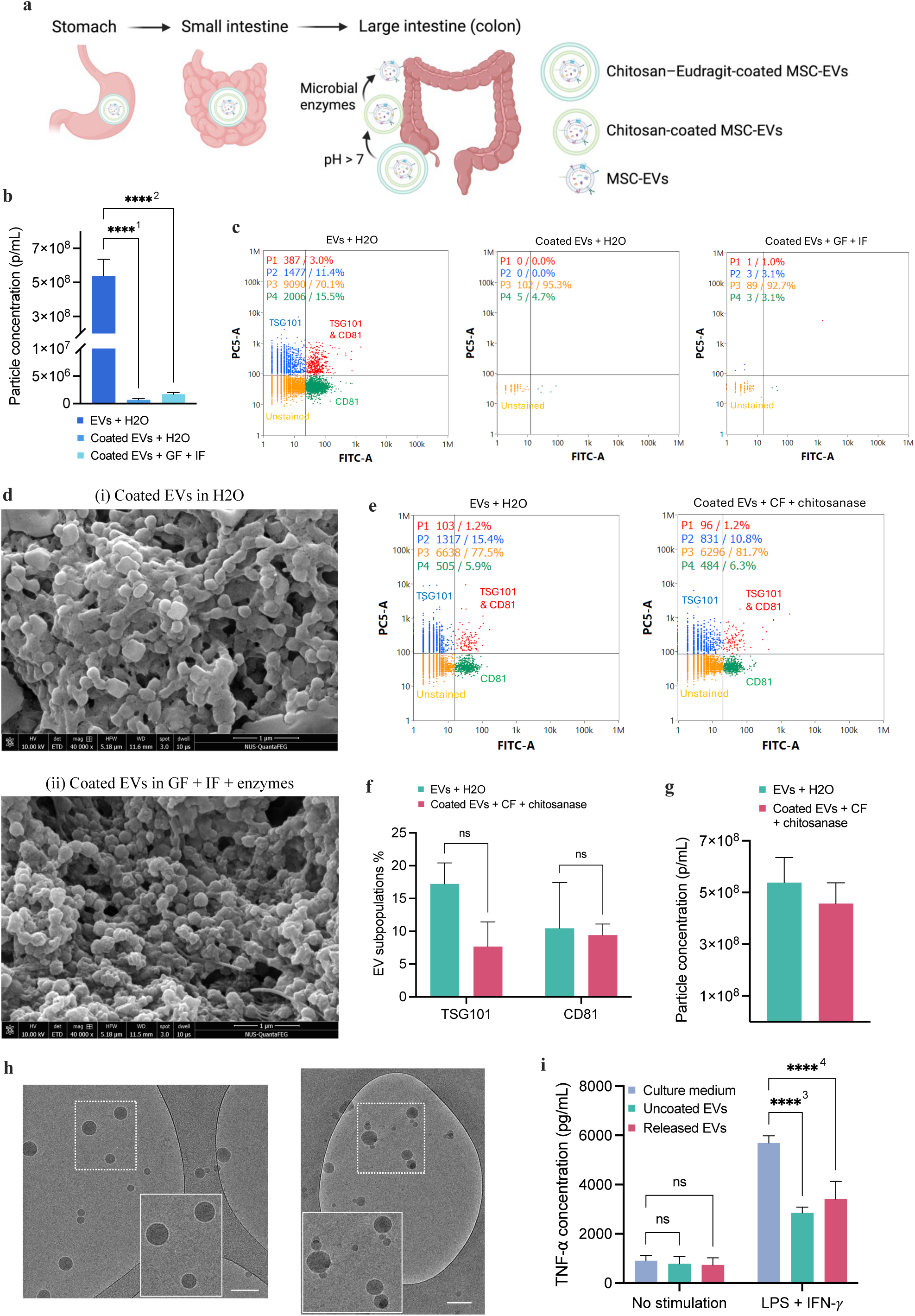
Stability of coated EVs in gastrointestinal environment and colon-targeted EV release. **(a)** Schematic representation of the stability and sequential release of coated EVs in the gastrointestinal (GI) tract. Coated MSC-EVs remain intact in the stomach and small intestine. Upon reaching the ileocolonic region, the outer Eudragit layer dissolves at pH > 7, and the inner chitosan layer is subsequently degraded by microbial enzymes in the colon, releasing MSC-EVs at the target site. Created using BioRender.com. **(b)** Particle concentration in the supernatant of coated EVs. Coated EVs both in water (pH 6.5) and after incubation with simulated gastric fluid (GF) and intestinal fluid (IF) showed significantly lower particle numbers in solution compared with uncoated EVs (^1^**** and ^2^**** p < 0.0001). The particle concentration for coated EVs was below the detection limit (1×10^8^ p/mL) for the flow nanoanalyzer (NanoFCM). **(c)** Representative NanoFCM dot plots showing percentage of EVs stained for TSG101 and/or CD81. In the supernatant of coated EVs both in water (pH 6.5) and GF + IF, only a small number of events were recorded and ≥ 90% of particles detected were unstained. **(d)** Scanning electron microscopy (SEM) images of coated EVs in (i) water (pH 6.5) and (ii) GF + IF with digestive enzymes pepsin and pancreatin. **(e)** Representative dot plots showing gated populations in stained uncoated EVs and coated EVs incubated with simulated colonic fluid (CF) and chitosanase based on fluorescence intensities, FITC-A: anti-CD81 (488) vs. PC5-A: anti-TSG101 (647), measured with flow nanoanalyzer (NanoFCM). P1: Double-positive for TSG101 and CD81; P2: TSG101-only positive; P3: Double-negative (unstained); P4: CD81-only positive. **(f)** Percentage of EV subpopulations positive for TSG101 and CD81 relative to total particles detected by nanoflow cytometry. The proportion of TSG101- and CD81-positive EVs released from coated EVs in CF + chitosanase were not significantly different to uncoated EVs (p > 0.05). **(g)** Particle concentration measured by nanoflow cytometry. 85% of particles were released from the coating within 6 h incubation with CF + chitosanase. **(h)** Cryogenic electron microscopy (cryo-EM) images of vesicles released from the coating formulation. Scale bar: 100 nm. **(i)** Effect of released EVs on TNF-⍺ levels in J774A.1 macrophages measured by ELISA. EVs released from the coating formulation did not activate unstimulated macrophages and reduced TNF-⍺ production in LPS/IFN-𝛾-activated macrophages to the same extent as uncoated EVs (^3^**** and ^4^**** p < 0.0001). Data are presented as mean ± SD (n = 3). **(b)** One-way ANOVA with post-hoc Dunnett’s test. **(i)** Two-way ANOVA with post-hoc Dunnett’s test. **(f)** Multiple paired t-tests. (**** p < 0.0001).

To assess stability in the upper GI tract, coated EVs were incubated with simulated gastric fluid (GF) followed by simulated intestinal fluid (IF). The supernatant was then analysed to determine whether EVs had been released from the coating. The particle concentration in GF and IF was below the detection limit for the NanoFCM flow nanoanalyzer (1×10^8^ particles/mL) and similar to background noise, indicating that the coating did not degrade and remained stable around the EVs (Figure 6b). Consistently, nanoflow cytometry scatter plots showed that of the few events (< 120) detected, ≥ 90% of particles were negative for TSG101 and/or CD81 (Figure 6c), suggesting that they may have been coating debris rather than EVs. SEM images obtained before and after incubation with GF and IF in the presence of digestive enzymes pepsin and pancreatin further confirmed that the coating remained intact and protected EVs from degradation, as individual particles were still visible within the coating matrix (Figure 6d and Supplementary Figure 10b).

To evaluate release of EVs in the colon, coated EVs were incubated with simulated colonic fluid (CF) and chitosanase to mimic enzymatic degradation of chitosan. Analysis by nanoflow cytometry revealed the presence of TSG101- and CD81-positive EVs (Figure 6e) at percentages comparable to uncoated EVs (Figure 6f). Particle concentration measurements showed that ∼85% of EVs were released from the coating after 6 h (Figure 6g) and cryo-EM images confirmed that the released EVs preserved their structural integrity (Figure 6h). EVs released from the coating formulation also retained their biological activity as evidenced by the reduction in TNF-⍺ levels in activated J774A.1 macrophages (Figure 6i).

These results demonstrate that the double-coating formulation remains intact under conditions simulating the stomach and small intestine, preventing premature release or degradation of EVs, while enabling rapid EV release in conditions mimicking the colon.

### Orally administered coated MSC-EVs alleviate disease severity in a DSS-induced colitis mouse model

We used a dextran sodium sulfate (DSS)-induced colitis mouse model to evaluate the therapeutic efficacy of coated MSC-EVs *in vivo*. Acute colitis was induced in male C57BL/6J mice by adding 3% DSS to drinking water for 5 days (Figure 7a). MSC-EVs (1×10^9^ particles/mouse) were administered on days 1–5 via intravenous (IV) injection and by oral gavage (uncoated EVs and coated EVs). Mice were monitored daily and clinical signs, including weight loss, diarrhoea and presence of blood in stools, were scored and summed to generate the disease activity index (DAI). On day 6, mice were euthanised and colons were collected for analysis.

**Figure 7.**
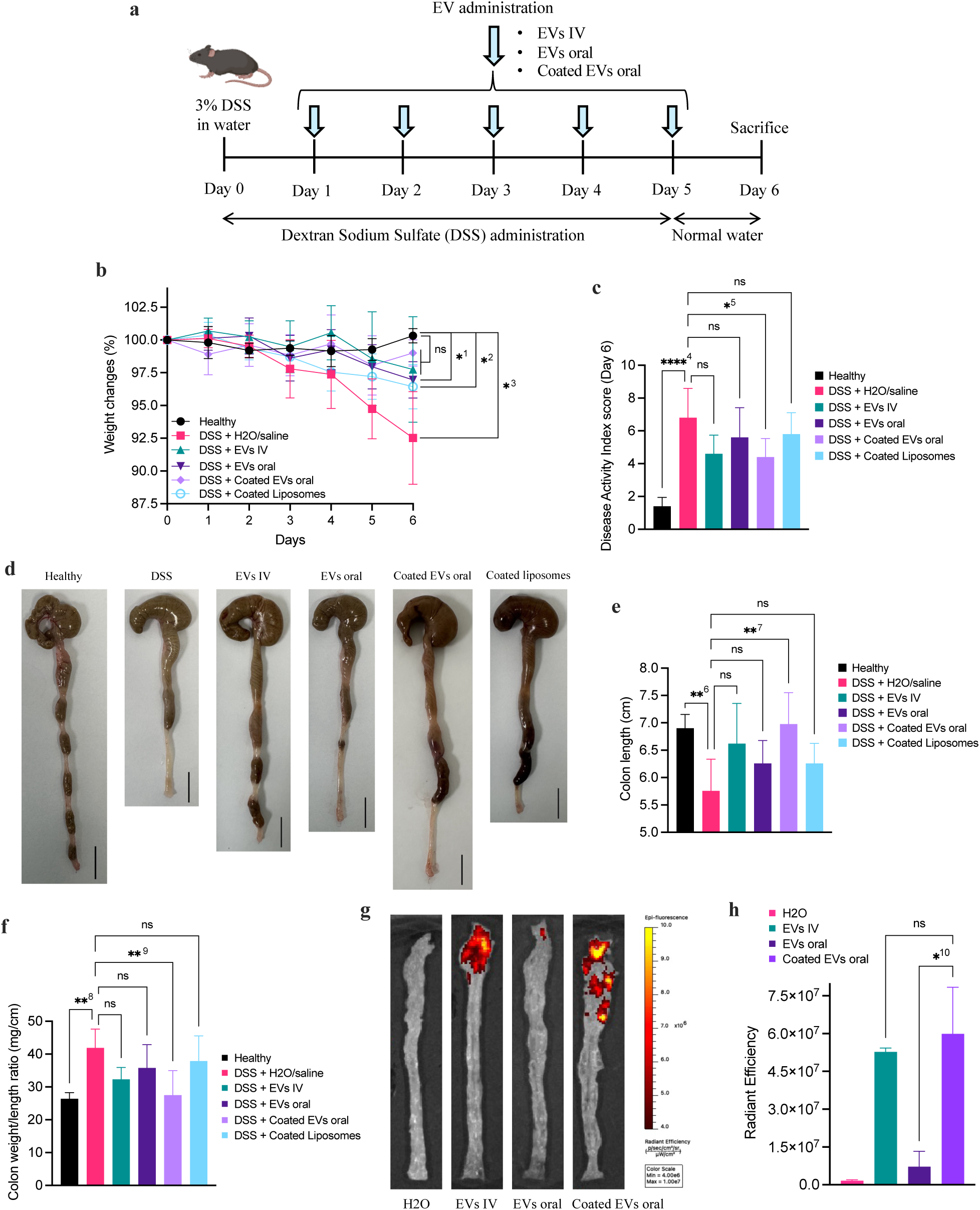
Therapeutic efficacy of coated MSC-EVs in a DSS-induced colitis mouse model. **(a)** Study design for dextran sodium sulfate (DSS)-induced colitis in mice and dosing schedule of MSC-EVs. 3% (w/v) DSS was administered in drinking water for 5 days to induce acute colitis in mice. MSC-EVs (1×10^9^ particles/mouse/day) were given on days 1–5 via intravenous (IV) injection (EVs IV) and by oral gavage for uncoated EVs (EVs oral) and coated EVs (coated EVs oral). **(b)** Weight changes in mice over 6 days. Mice treated with EVs via IV injection and coated EVs administered orally showed minimal changes in body weight, similar to healthy mice. There was significant body weight loss in mice treated with uncoated EVs orally (^1^* p = 0.0122), coated liposomes (^2^* p = 0.0164) and in DSS control mice (^3^* p = 0.0231). **(c)** Disease activity index (DIA) scores on day 6. DAI combines scores for weight loss, stool consistency and faecal occult blood. The DIA score was the highest for DSS control mice and the lowest for healthy mice (^4^**** p < 0.0001). Mice treated with coated EVs orally were the only treatment group that showed a significantly lower DAI score (^5^* p = 0.0407). (EVs IV p = 0.0670). **(d)** Photographs of colons on day 6. Scale bar 1 cm. **(e)** Colon length measurements. Treatment with coated EVs significantly increased the colon length of DSS mice (^7^** p = 0.0041) and showed similar results as healthy colons (^6^** p = 0.0075). (EVs IV p = 0.0539). **(f)** Colon weight to length ratio. Mice treated with coated EVs showed a significant reduction in the colon weight/length ratio (^9^** p = 0.0038) to same extent as healthy colons (^8^** p = 0.0019). (EVs IV p = 0.0705). Data are presented as mean ± SD (n = 5). **(g)** Biodistribution of MSC-EVs in the colon of DSS mice 24 h post dosing. WGA-AF680–labelled EVs were administered via IV injection and oral gavage for uncoated and coated EVs and the fluorescence in the colon was imaged using IVIS (Revvity). **(h)** Quantification of EV fluorescence signal (radiant efficiency) in the colon 24 h after administration. Coating of EVs significantly increased fluorescence intensity in the colon compared with uncoated EVs administered orally (^10^* p = 0.0131). Data are presented as mean ± SD (n = 2). **(b)** Two-way ANOVA with Dunnett’s test. **(c)**, **(e)**, **(f)** and **(h)** One-way ANOVA with Dunnett’s test. (* p < 0.05, ** p < 0.01, **** p < 0.0001).

DSS treatment resulted in significant body weight loss after 6 days, an effect that was also observed in mice given uncoated MSC-EVs orally (Figure 7b). By contrast, mice that received coated EVs orally and EVs via IV injection maintained their body weight throughout the study, similar to healthy controls. However, only coated EVs significantly reduced DAI scores on day 6 compared with other EV treatment groups, indicating a stronger reduction in colitis severity (Figure 7c). At necropsy, the colons of each group were photographed (Figure 7d). DSS exposure shortens and thickens the colon as a result of inflammation and damage to the intestinal mucosa. Treatment with coated EVs significantly increased colon length (Figure 7e) and reduced colon weight/length ratio (Figure 7f), normalising these parameters to values observed in healthy controls. By comparison, uncoated EVs administered orally produced outcomes similar to the DSS control group, emphasising the requirement for formulation to preserve the biological activity of MSC-EVs. Additionally, coated EVs induced a stronger therapeutic response than intravenous EVs administered at the same relatively low dose, suggesting that oral delivery may be more effective when the site of action is localised to the colon. The limited efficacy of intravenous EVs is likely due to their rapid clearance from the circulation and insufficient targeting or retention at the site of inflammation. The enhanced efficacy of coated EVs may also reflect a synergistic contribution from chitosan, which has intrinsic anti-inflammatory properties [46]. To confirm that the therapeutic effect of coated EVs was predominantly mediated by EVs rather than the coating, we included an additional group treated with coated liposomes. These mice exhibited responses similar to the DSS control group, including weight loss, elevated DAI scores, and colon shortening and thickening, demonstrating that the coating alone did not confer therapeutic benefit in DSS-induced colitis.

We also investigated the biodistribution of MSC-EVs in DSS mice. The double-coating formulation significantly increased EV accumulation in the colon 24 h after oral administration compared with uncoated EVs (Figure 7g and 7h). Coated EVs also showed significantly lower liver localisation (Supplementary Figure 12), suggesting reduced systemic uptake and translocation to the liver.

Overall, our study demonstrated that MSC-EVs are disintegrated in the gastrointestinal environment, resulting in a loss of biological activity. To facilitate oral delivery, we developed a chitosan–Eudragit double-coating formulation that improves EV stability in GI fluids and targets delivery to the colon. This coating system protected MSC-EVs from degradation by digestive conditions and enabled the release of structurally intact, biologically active vesicles in simulated colonic fluid. The *in vivo* data suggest that functional EVs were released in the inflamed colon, as orally administered coated EVs elicited a therapeutic response not observed with uncoated EVs. Our preliminary *in vivo* studies also showed that orally administered coated EVs exerted a stronger therapeutic effect than EVs delivered intravenously. Additional biodistribution and pharmacokinetic studies would be needed to determine whether this enhanced efficacy is due to improved targeting, greater retention, or both, within inflamed intestinal tissues.

Although MSC-EVs have been extensively studied in colitis models, oral delivery remains relatively under-explored. Prior to this work, only two studies (Gan et al. [47] and Deng et al. [48]) had reported the therapeutic efficacy of orally administered MSC-EVs in colitis. Our findings therefore contribute to the emerging evidence supporting oral delivery of MSC-EVs as a promising therapeutic approach for IBD.

Our *in vivo* evaluation was restricted to macroscopic endpoints, including disease activity index and colon morphology, which offer only a partial view of therapeutic efficacy. More comprehensive characterisation, such as histological assessment and cytokine profiling, is required to substantiate these findings and determine the extent of mucosal healing and inflammation resolution.

## Conclusion

MSC-EVs possess strong therapeutic potential for IBD owing to their regenerative and immunomodulatory properties. Effective treatment of IBD requires localised drug delivery to the intestine, making oral administration an attractive and patient-friendly route. In this study, we demonstrated that MSC-EVs are unstable in gastrointestinal fluids, underscoring the need for an optimised delivery system to enable oral administration. We developed a double-coating formulation that protects MSC-EVs from degradation in the GI environment and targets delivery to the colon. Our preliminary *in vivo* findings suggest that, with suitable formulation, oral delivery of MSC-EVs may offer a promising strategy for treating intestinal inflammation.

## Methods

### Cell culture

Human bone marrow mesenchymal stem cells (hBM-MSCs) were purchased from Lonza (product #PT-2501) in 2022 and 2024, certified mycoplasma-free, and used between passages 3–5. The primary cells were cultured in Dulbecco’s modified eagle medium (DMEM) containing 1 g/L glucose (product #31885-023, Thermo Fisher Scientific) supplemented with 10% (v/v) EV-depleted foetal bovine serum (FBS) (product #F9665, Sigma-Aldrich) and 1% (v/v) antibiotic-antimycotic solution 100X containing penicillin, streptomycin and amphotericin B (product #15240-062, Thermo Fisher Scientific). EV-depleted FBS was prepared using the gold standard method [49] by subjecting FBS to ultracentrifugation (Optima^TM^ XPN, Beckman Coulter) at 100,000 𝑔 in a fixed-angle rotor (45 Ti, Beckman Coulter) for 18 h at 4 °C. The FBS supernatant was then collected and sterile filtered using 0.22 μm filters for use in cell culture.

Human epithelial colorectal adenocarcinoma (Caco-2; RRID:CVCL_0025) and murine J774A.1 macrophage (RRID:CVCL_0358) were purchased from the European Collection of Authenticated Cell Cultures in 2020 and 2021, respectively, and certified mycoplasma-free. Caco-2 cells were used between passages 25–50 and J774A.1 cells were used between passages 10–30. Both cell lines were cultured in DMEM containing 4.5 g/L glucose (product #41966-029, Thermo Fisher Scientific) supplemented with 10% (v/v) FBS, 1% (v/v) non-essential amino acids (NEAA) (product #25-025-Cl, Corning) and 1% (v/v) antibiotic-antimycotic solution 100X containing penicillin, streptomycin and amphotericin B.

All cells were maintained at 37 °C in a humidified incubator containing 5% CO^2^.

### Isolation of MSC-EVs

EVs were isolated using a one-step sucrose cushion ultracentrifugation technique, as described in detail [50] (Supplementary Figure 1). The conditioned medium (CM) of hBM-MSCs was harvested and pre-cleared of dead cells and cellular debris through several rounds of differential centrifugation: 500 𝑔 for 5 min at 4 °C twice, then 2000 𝑔 for 15 min at 4 °C; followed by filtration of the supernatant through 0.22 μm sterile filter. Pre-cleared CM was either stored at −80 °C or directly processed for EV isolation using the Optima^TM^ XPN ultracentrifuge (Beckman Coulter). Thickwall polycarbonate ultracentrifuge tubes (product #355631, Beckman Coulter) were filled with 22.5 mL of pre-cleared CM, then 3 mL of 25% (w/w) sucrose solution (product #S/8600/60, Fisher Scientific) prepared in deuterium oxide (D_2_O) (product #151882, Sigma Aldrich) was slowly layered below the CM using a glass Pasteur pipette. The ultracentrifuge tubes were placed in a swing-out rotor (SW32 Ti or SW28 Ti, Beckman Coulter) and subjected to ultracentrifugation at 100,000 𝑔 for 1.5 h at 4 °C. The sucrose solution was then collected (2 ml per tube) and added to filtered PBS in polycarbonate ultracentrifuge tubes (product #355622 or # 355618, Beckman Coulter) as a washing step for EV purification. The ultracentrifuge tubes were subjected to ultracentrifugation in a fixed-angle rotor (45 Ti or 70 Ti, Beckman Coulter) at 100,000 𝑔 for 1.5 h at 4 °C. The supernatant was then discarded and the pellet of EVs was resuspended in filtered PBS and stored at 4 °C (short term) or −80 °C (long term).

### Characterisation of MSC-EVs

#### Nanoparticle tracking analysis (NTA)

The size distribution and concentration of EVs was measured by NTA using a NanoSight instrument (LM10 or NS300, Malvern Panalytical). EV samples were diluted in filtered PBS to obtain 20-80 particles in the viewing frame for optimal tracking. The samples were injected into the laser viewing module and the camera level was adjusted to 14-15, with screen gain 1. The analysis detection threshold was set at 5-7. For each sample, video recordings of 30-60 seconds duration were captured and analysed using the NanoSight NTA software. A size distribution histogram was generated based on the Brownian motion of the particles in each sample.

#### Bicinchoninic acid (BCA) assay

The total protein concentration of EVs was quantified using the QuantiPro^TM^ BCA assay kit (Sigma-Aldrich). EV samples and varying concentrations (0-20 µg/mL) of bovine serum albumin (BSA) standards prepared in PBS were added to a 96-well plate (50 µL/well). The working reagent (prepared according to the manufacturer’s instructions) was then added to the wells in equal volume (50 µL/well) and the plate was incubated at 60 °C for 1 hour. The absorbance was measured at 562 nm using a plate reader (Infinite 200 Pro, Tecan) and a BSA standard curve was used to determine the protein concentration of EV samples.

#### Zeta potential

The zeta potential of EVs was measured at 25 °C using a Zetasizer instrument (Zetasizer Nano or Ultra, Malvern Panalytical) and analysed using the ZS Explorer software. EV samples were diluted in PBS (minimum concentration 1×10^11^ particles/ml) prior to loading the samples in folded capillary zeta cells (DTS1070, Malvern Panalytical).

#### Cryogenic electron microscopy (cryo-EM)

3.5 µl of EV samples (minimum concentration 1×10^9^ particles/mL) was applied twice to a Tedpella lacey fomvar/carbon copper grid (Ted Pella, Inc.) that was glow-discharged in air for 60 seconds. Then the grid was blotted for 1.5 seconds (blot force 1) at 22 °C and 100% humidity, followed by a plunge into liquid ethane using FEI Vitrobot Mark IV. The micrographs were recorded using a 200 kV Tecnai Arctica cryo-transmission electron microscope equipped with a Falcon 3EC direct electron detector. Images were collected at a magnification of 39,000X and 53,000X, yielding the pixel sizes of 2.79 and 2.01 Ångstroms per pixel (Å/px), respectively.

#### Exosome antibody array kit

The presence of EV markers was detected using the Exo-Check^TM^ Exosome Antibody Array kit (System BioScience) following the manufacturer’s protocol. The array had eight positive markers (tetraspanins CD63 and CD81, EpCAM: epithelial cell adhesion molecule, ANXA5: annexin A5, TSG101: tumor susceptibility gene 101, FLOT1: flotillin-1, ICAM: intercellular adhesion molecule 1, ALIX: programmed cell death 6 interacting protein) and four controls including cis-Golgi matrix protein (GM130) as a negative marker for cellular contamination during exosome isolation, two positive controls and a blank control.

#### Super-resolution microscopy

The EV Profiler 2 kit (Oxford Nanoimaging) was used for visualising and phenotyping single EVs. EVs were stained with a tetraspanin antibody panel consisting of anti-CD63 (561), anti-CD81 (647) and anti-CD9 (488). Samples were processed according to manufacturer’s instructions to immobilise the stained EVs on the assay chips. Three fields of view were recorded for each sample and super-resolution images were acquired using direct stochastic optical reconstruction microscopy (dSTORM) on a Nanoimager instrument (Oxford Nanoimaging). EV clusters were filtered based on circularity, density of localizations and area to remove background and non-EV-related clusters. Colocalizations of the different tetraspanins were analysed using the CODI platform (Oxford Nanoimaging).

### Cell proliferation assay

The MTS assay was used to assess cell viability, metabolic activity and proliferation. Caco-2 cells were seeded at different densities (5000-150,000 cells/well) in 48-well plates (Thermo Fisher Scientific) and incubated overnight (12 h) at 37 °C to allow cells to adhere to the wells. The next day, the cells were washed with PBS before 200 µL of serum-free medium and 20 µL of MTS reagent (CellTiter 96^®^ AQ_ueous_ One Solution, Promega) were added to the wells. Cells with active mitochondrial metabolism convert the MTS tetrazolium reagent into a purple-coloured formazan product that is soluble in cell culture medium and can be quantified by measuring absorbance at 490 nm [51]. After 2 h incubation at 37 °C, the absorbance of the formazan product generated in the wells was measured at 490 nm using an Infinite 200 Pro plate reader (Tecan). The absorbance of formazan was directly proportional to the number of viable Caco-2 cells and a standard curve was constructed to plot the absorbance at 490 nm against different Caco-2 cell densities per well (Supplementary Figure 2).

For the cell proliferation assay, Caco-2 cells were seeded in 48-well plates (Thermo Fisher Scientific) at a density of 1.5×10^5^ cells/mL (200 μL/well) and incubated at 37 °C for 24 h. The next day, culture medium was removed from the wells and Caco-2 cells were incubated with increasing concentrations (10^7^, 10^9^ and 10^11^ particles/mL) of MSC-EVs prepared in serum-free culture medium (200 μL/well). Serum-free cell culture medium was used as a control and 10% (v/v) dimethyl sulfoxide (DMSO) (product #D2438, Sigma Aldrich), also in serum-free culture medium, was used as a negative control for cell proliferation since it is toxic to cells. The MTS assay was performed daily for 4 consecutive days. The sample and control solutions were discarded from the wells and the cells were washed with PBS, then 200 μL of serum-free medium was added to each well followed by 20 μL of MTS reagent. The plate was incubated for 2 h at 37 °C and the absorbance at 490 nm was measured using an Infinite 200 Pro plate reader (Tecan). Caco-2 cell density/well was determined using the constructed standard curve.

### Wound healing assay

Culture-inserts (Ibidi) consisting of 2 wells separated by a wall were placed in 12-well plates (Thermo Fisher Scientific). Caco-2 cells were seeded in each well of the culture-insert at a density of 7×10^5^ cells/mL (70 µL/well) and incubated at 37 °C for 24 h. The following day, culture-inserts were removed using sterile tweezers, creating a cell-free 500 µm gap between the two cell layers, and the wells were washed with PBS to remove cell debris. Caco-2 cells were then incubated with 1 mL of serum-free culture medium containing increasing concentrations (10^7^, 10^9^ and 10^11^ particles/mL) of MSC-EVs. Serum-free cell culture medium was used as a control. Images of the wounds were captured at regular intervals (0 h, 24 h, 48 h and 72 h) using an Eclipse Ts2R inverted microscope (Nikon) on a 10X objective. The images were then processed using a *wound_healing_size_tool* plugin [52] for ImageJ to measure the wound area and average wound width. Wound healing was quantified using two metrics: (1) percentage of area reduction or wound closure and (2) rate of cell migration (R_M_) [39].

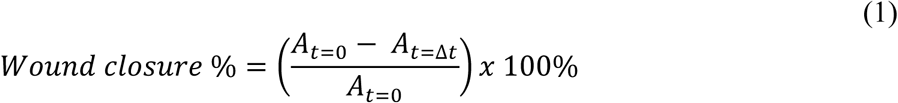

Where A_t = 0_ is the initial wound area and A_t = Δt_ is the wound area after *n* h, both in μm^2^.

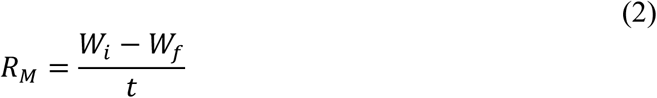

R_M_ is the rate of cell migration (µm/h). W_i_ is the initial wound width, W_f_ is the final wound width, both in μm, and t is the duration of the migration in h.

### Pro-inflammatory cytokine assay

J774A.1 macrophages were seeded in 48-well plates (Thermo Fisher Scientific) at a density of 1×10^5^ cells/mL (0.4 mL/well) in culture medium containing 100 ng/mL lipopolysaccharide (LPS) from Escherichia coli O111:B4 (product #L2630, Sigma Aldrich) and 25 ng/mL interferon-𝛾 (IFN-𝛾) (product #GFH77, Cell Guidance Systems) to induce inflammation. The cells were incubated at 37 °C for 24 h. The next day, activated macrophages were treated with increasing concentrations of MSC-EVs (final concentration in wells: 10^7^, 10^9^ and 10^11^ particles/mL) and incubated for a further 24 h. 1 µg/mL Dexamethasone (product #D4902, Sigma Aldrich) was used as a positive control for anti-inflammatory activity and cell culture medium was used as a negative control. Cytokines (TNF-α and IL-6) released from J774A.1 macrophages were quantified using a Mouse Luminex^®^ Discovery assay (product #LXSAMSM, Bio-Techne), following the manufacturer’s protocol. The median fluorescence intensity was measured using a Luminex^®^ Flexmap3D^®^ Reader and standard curves for each cytokine were generated to quantify cytokine levels in the samples.

### Oxidative stress assay

J774A.1 macrophages were seeded in 24-well plates (Thermo Fisher Scientific) at a density of 2×10^5^ cells/mL in culture medium without antibiotics (0.5 mL/well) containing 100 ng/mL LPS and 100 nM phorbol 12-myristate 13-acetate (PMA; product #P8139, Sigma Aldrich) to induce oxidative stress. The cells were incubated at 37 °C for 24 h. The next day, activated macrophages were treated with increasing concentrations of MSC-EVs (final concentration in wells: 10^7^, 10^9^ and 10^11^ particles/mL) and incubated for a further 24 h. Cellular oxidative stress was quantified by measuring total reactive oxygen species (ROS) using 2’,7’-dichlorodihydrofluorescein diacetate (DCFH-DA; product #D6883, Sigma Aldrich). DCFH-DA is taken up by cells where cellular esterase cleaves off the acetyl groups, resulting in DCFH, which is then oxidised by ROS and converted to DCF, which emits green fluorescence that can be measured [53]. J774A.1 macrophages were washed with DMEM, then stained with 500 µL of DCFH-DA solution (50 µM) and incubated at 37 °C for 30 min. The cells were subsequently washed with DMEM and incubated with 200 µL RIPA buffer (product #R0278, Sigma Aldrich) on ice for 5 min. The lysate was collected and centrifuged at 16000 *g* for 10 min at 4 °C. The supernatant was collected and the fluorescence (Ex/Em 485/530) was measured using an Infinite 200 Pro plate reader (Tecan). 1 mmol/L Nω-Nitro-L-arginine methyl ester (L-NAME; product #N5751, Sigma Aldrich) was used as a positive control for antioxidant activity as it indirectly reduces cellular ROS production by preventing peroxynitrite formation through the inhibition of the enzyme nitric oxide synthase, which produces nitric oxide (NO). Cell culture medium without antibiotics was used as a negative control.

### Transport and uptake of MSC-EVs in vitro

#### Labelling of EVs

MSC-EVs were labelled with fluorescent Wheat Germ Agglutinin (WGA) conjugates. Briefly, MSC-EVs were incubated with 0.1 mg/mL WGA-Alexa Fluor 488 (WGA-AF488; product #W11261, Thermo Fisher Scientifc) or WGA-Alexa Fluor 680 (WGA-AF680; product #W32465, Thermo Fisher Scientifc) at 37 °C for 30 min under gentle agitation, protected from light (Supplementary Figure 3a). The labelled EVs were purified using Exo-spin columns (EX03 Exo-spin™ mini, Cell Guidance Systems) and the recovery of EVs was determined using nanoparticle tracking analysis (NTA) (Supplementary Figure 3b). Prior to each assay, labelled EVs were analysed by NTA to measure particle size and concentration for accurate *in vitro* dosing. For permeability studies, a fluorescence standard curve was also constructed to determine the particle concentration in samples.

#### Transport of EVs across intestinal epithelial cells

The intestinal permeability of MSC-EVs was determined using an established model of the intestinal epithelium. Caco-2 colon epithelial cells were seeded at a density of 1×10^5^ cells/cm^2^ on Transwell^®^ cell culture inserts (polycarbonate filter, 1.12 cm^2^ area, 0.4 μm pore size, 1 mL in apical compartment and 1.5 mL in basolateral compartment; Corning). The Caco-2 cells were maintained in culture on a semipermeable membrane insert for 19-21 days until they differentiated into enterocyte-like cells of the small intestine [54]. The transepithelial electrical resistance (TEER) was measured every 2-3 days of the culture using an Epithelial Volt Ohm Meter (EVOM) (World Precision Instruments) to assess the development of epithelial barrier integrity and formation of tight junctions in Caco-2 monolayers (Supplementary Figure 4). The STX2/chopstick electrodes of the EVOM were sterilised in 70% ethanol and neutralised in DMEM prior to taking measurements. The resistance measurements were corrected for the blank and multiplied by the Transwell^®^ insert filter area (1.12 cm^2^) to obtain TEER values in ohm per cm^2^ (Ω·cm^2^). TEER values ≥ 500 Ω·cm^2^ are indicative of fully formed tight junctions in the Caco-2 monolayer [55].

For the permeability assay, the culture medium was removed from differentiated Caco-2 cells in transwells and the cells were incubated with Hanks’ Balanced Salt Solution (HBSS) (product #H8264, Sigma-Aldrich) at 37 °C for 1 hour. The HBSS solution in the apical (or upper) compartment was then removed and MSC-EVs (1×10^9^ particles/mL) labelled with WGA-AF488 (prepared in HBSS) were applied to the Caco-2 monolayer and incubated at 37 °C for 3 h. Samples (100 µL) were collected from the basolateral (or lower) compartment every 30 min and replaced with an equal volume of HBSS to maintain equilibrium. The particle concentration of transported EVs was measured using fluorescence intensity at Ex/Em 490/530 and nanoparticle tracking analysis (NTA) (Supplementary Figure 5a). Hourly samples were also analysed for particle size using NTA (Supplementary Figure 5b) and cryogenic electron microscopy (cryo-EM) was used to image transported vesicles (Supplementary Figure 5c). The TEER was measured before and after the permeability study to monitor changes in the Caco-2 barrier integrity (Supplementary Figure 5d).

#### Cellular uptake of EVs

Caco-2 epithelial cells were seeded at a density of 1×10^5^ cells/cm^2^ on Transwell^®^ cell culture inserts (polycarbonate semipermeable filter, 1.12 cm^2^ area, 0.4 μm pore size; Corning) and maintained in culture for 19-21 days until differentiation. J774A.1 macrophages were seeded in 12-well plates (Thermo Fisher Scientific) at a density of 5×10^5^ cells/mL (1 mL/well). MSC-EVs labelled with WGA-AF488 (1×10^9^ particles/mL) were applied to the cells and incubated at 37 °C for 4 h. Subsequently, differentiated Caco-2 cells were detached using TrypLE^TM^ Express (1X) stable trypsin replacement enzyme (product #12605-010, Thermo Fisher Scientific) and J774A.1 macrophages were detached using a cell scraper and centrifuged at 1500 RPM for 10 min. The cells were then stained with DAPI in 2% bovine serum albumin (BSA) and analysed with a Cytoflex LX flow cytometer (Beckman Coulter) at FITC channel to quantify EV uptake in differentiated Caco-2 intestinal epithelial cells (Supplementary Figure 6a) and J774A.1 macrophages (Supplementary Figure 7a). Unstained cells were used as a control to set the gating (Supplementary Figure 8). At least 10,000 events were analysed and data were processed using CytExpert 2.5 software (Beckman Coulter).

The uptake of MSC-EVs in Caco-2 intestinal epithelial cells was imaged using confocal microscopy (Supplementary Figure 6b). Differentiated Caco-2 cells on Transwell^®^ cell culture inserts with polyester filter (Corning) were incubated with WGA-AF488–labelled EVs (1×10^10^ particles/mL) at 37 °C for 4 h. The cells were washed with PBS and fixed with paraformaldehyde 4% fixative in PBS (bioWorld) for 20 min at room temperature. The cells were then permeabilised with 0.2% (v/v) Triton X-100 (Sigma Aldrich) in PBS for 5 min and blocked against non-specific binding with 5% (w/v) blocking buffer (skimmed milk powder in PBS) and 0.05% (v/v) Triton X-100 for 30 min at room temperature. Afterwards, the cells were incubated with rabbit anti-ZO-1 primary polyclonal antibody (product #40-2200, Thermo Fisher Scientific) (1:150) in 1% blocking buffer and 0.5% (v/v) Triton X overnight at 4 °C. After washing with PBS the next day, the cells were incubated with chicken anti-rabbit IgG (H+L) cross-adsorbed secondary antibody Alexa Fluor 647 (product #A21443, Thermo Fisher Scientific) (1:500) in 1% (w/v) blocking buffer and 0.05% (v/v) Triton X for 1 hour at room temperature protected from light to stain the tight junction protein zonula occludens (ZO)-1. Following another washing step, 1 drop of DAPI-containing mounting medium was added to the insert for 10 min to stain the cell nuclei. The insert membrane was then excised using a scalpel and laid flat onto a microscopy µ-slide (Ibidi) using 0.5% (w/v) agarose in PBS. Fluorescence images were captured using a Nikon Eclipse Ti Inverted Spinning Disk confocal microscope with a Yokogawa CSU-X1 spinning disk unit and Andor EMMCD Camera. Images were processed using Fiji software.

The uptake of MSC-EVs in J774A.1 macrophages was imaged using imaging flow cytometry (Supplementary Figure 7b). J774A.1 macrophages were seeded in 100 mm cell culture dishes (Thermo Fisher Scientific) at a density of 1×10^6^ cells/mL (5 mL/dish) and incubated with WGA-AF680–labelled EVs (1×10^9^ particles/mL) at 37 °C for 4 h. The cells were manually detached using a cell scraper, centrifuged at 1500 RPM for 10 min and re-suspended in 500 µL PBS for analysis using the Cytek Amnis ImageStream Mark II imaging flow cytometer.

### MSC-EVs in a co-culture model of intestinal inflammation

A Caco-2/J774A.1 co-culture that mimics intestinal inflammation *in vitro* was established as described in detail [43]. Caco-2 cells were seeded at a density of 1×10^5^ cells/cm^2^ on Transwell^®^ cell culture inserts (polycarbonate filter, 1.12 cm^2^ area, 0.4 μm pore size, 1 mL in apical compartment and 1.5 mL in basolateral compartment; Corning). Caco-2 cells were maintained in culture for 19–21 days until they differentiated into polarised intestinal epithelial monolayers. Stabilisation of transepithelial electrical resistance (TEER) values (∼2000 Ω·cm²) indicated the presence of an intact epithelial barrier with fully formed tight junctions. Differentiated Caco-2 cells were primed with an optimised cytokine cocktail of TNF-α, IFN-γ and IL-1β (25 ng/mL) added to the basolateral compartment. On the same day, J774A.1 macrophages were seeded in 12-well plates (Corning) at a density of 5×10^4^ cells/well (1.5 mL/well) in culture medium containing LPS (100 ng/mL) and IFN-𝛾 (25 ng/mL). After 24 h, the two cell lines were combined into a co-culture with differentiated Caco-2 epithelial cells in the apical compartment and J774A.1 macrophages in the basolateral compartment to mimic the inflamed intestine. MSC-EVs were then applied to the inflamed Caco-2/J774A.1 co-culture either in the apical or basolateral compartment (final concentration 1×10^9^ particles/mL). The volumes in the apical and basolateral compartments of the Transwell^®^ cell culture system were maintained at equal level to prevent changes in osmolarity. Samples were collected from the basolateral compartment after 6 and 12 h to quantify pro-inflammatory cytokine (TNF-α and IL-6) production from macrophages using the Luminex^®^ assay. The TEER was measured at 12 and 16 h to monitor changes in the epithelial barrier integrity. TEER values were normalised to the baseline values of Caco-2 monocultures and expressed as percentages. A healthy co-culture of differentiated Caco-2 cells and J774A.1 macrophages was used as a control.

### Stability of MSC-EVs in gastrointestinal fluids

MSC-EVs were incubated with dissolution media that simulate human gut fluids to mimic gastrointestinal digestion *in vitro*. EVs (∼2×10^9^ particles/mL) were first incubated with fasted state simulated gastric fluid (FaSSGF; pH 1.7), prepared according to the manufacturer’s instructions using FaSSGF buffer concentrate (product #FASGBUF, Biorelevant) and 3F Powder (product #FFF02, Biorelevant), in a 1:1 (v/v) ratio for 2 h at 37 °C with gentle orbital shaking. The resulting EV mixture was then incubated with fasted state simulated intestinal fluid (FaSSIF; pH 6.6), prepared according to the manufacturer’s instructions using FaSSIF buffer concentrate (product #FASBUF, Biorelevant) and 3F Powder, in a 1:1 (v/v) ratio for another 2 h at 37 °C with gentle orbital shaking. Digestive enzymes were also included following the static *in vitro* digestion protocol [56]. Pepsin (product #P7012, Sigma Aldrich) was added to simulated gastric fluid (GF) at a final concentration of 2000 U/mL and pancreatin (product #P1750, Sigma Aldrich) was added to simulated intestinal fluid (IF) at a final concentration of 100 U/mL trypsin activity (based on TAME assay; 1 TAME unit = 19.2 USP units [57]). Milli-Q water and 10% sodium dodecyl sulfate (SDS) were used as controls for EV disintegration.

Following incubation with GF and IF in the presence of digestive enzymes, EVs were filtered through Exo-spin columns (EX03 Exo-spin™ mini, Cell Guidance Systems) and analysed using nanoparticle tracking analysis (NTA; Nanosight Pro NS500, Malvern Panalytical) to measure particle size distribution. MSC-EVs were also imaged using cryogenic electron microscopy (cryo-EM) 30 min and 1 hour after digestion to capture vesicle morphology. EVs incubated with GF and IF (without digestive enzymes) were stained with anti-CD81 488 (1:100000; product #CL488-65195, Proteintech) and anti-TSG101 647 (1:100000; product #CL647-67381, Proteintech) and analysed with a flow nanoanalyzer (NanoFCM) to determine particle concentration and percentage of EV subpopulations positive for transmembrane protein CD81 and/or cytosolic protein TSG101.

### Oral formulation of MSC-EVs

#### Double-coating formulation and characterisation

MSC-EVs were sequentially coated with chitosan followed by Eudragit S-100 to form chitosan–Eudragit-coated EVs, referred to as coated EVs (Supplementary Figure 9). A 0.1% (w/v) chitosan solution (product #448869, Sigma Aldrich) was prepared in 0.5% (v/v) acetic acid (pre-adjusted to pH 5.5–5.7 with NaOH; product #A113-50, Fisher Scientific). The chitosan solution was stirred at 500 RPM for 1–2 h at room temperature, then stored at 4 °C. MSC-EVs in PBS (10^9^–10^10^ particles/mL) were slowly added dropwise to an equal volume of 0.1% chitosan solution under magnetic stirring (500 RPM) at room temperature. Chitosan-coated EVs were purified using a Float-A-Lyzer dialysis device (1 mL, 1000 kDa MWCO; product #G235037, Repligen) to remove excess chitosan. Dialysis was performed at 4 °C with the membrane submerged in 50 mM sodium acetate buffer (pH 5.5–5.7) in a 500 mL beaker for 6 h with gentle stirring (100 RPM), replacing the buffer every 2 h. The chitosan-coated EVs were then characterized for particle size and concentration by nanoparticle tracking analysis (NTA), surface charge by zeta potential, and morphology by cryogenic electron microscopy (cryo-EM). A 0.2 % (w/v) Eudragit^®^ S-100 solution (Evonik) prepared in PBS (pH > 7) was slowly added dropwise to the chitosan-coated EVs under magnetic stirring (500 RPM) at room temperature. The appearance of white precipitates in the solution signalled the formation of a solid double-layer coating around the EVs. The addition of Eudragit was stopped when no further precipitates were observed and before the solution became visibly turbid due to excess polymer. The resulting chitosan–Eudragit-coated EVs were then washed several times with Milli-Q water (pH 6.5) by removing the supernatant and resuspending the precipitates in fresh water. A clear solution indicated removal of residual Eudragit not incorporated into the coating. The coated EVs were then stored at 4 °C until further use. Coated EVs were dried and imaged by scanning electron microscopy (SEM; JSM-6701F, Jeol).

#### Stability of coated EVs in gastrointestinal fluids

Coated MSC-EVs were first incubated with FaSSGF for 2 h (1:1 volume ratio), followed by FaSSIF for another 2 h (1:1 volume ratio) at 37 °C with gentle orbital shaking. The supernatant of coated EVs was then stained with anti-CD81 488 (1:100000) and anti-TSG101 647 (1:100000) and analysed with a flow nanoanalyzer (NanoFCM) to determine particle concentration of released EVs from the coating and percentage of CD81- and/or TSG101-positive EVs. Coated EVs were also incubated with FaSSGF containing pepsin (2000 U/mL) and FaSSIF containing pancreatin (100 U/mL trypsin activity) for 2 h at 37 °C with gentle orbital shaking. The coated EVs were then dried and visualised with scanning electron microscopy (SEM; JSM-6701F, Jeol). Uncoated EVs (∼2×10^9^ particles/mL) and coated EVs in water were used as controls.

#### EV release from double-coating formulation in vitro

Coated MSC-EVs were incubated with fasted state simulated colonic fluid (FaSSCoF; pH 7.8), prepared according to the manufacturer’s instructions using FaSSCoF Powder (product # COFAS01, Biorelevant), containing chitosanase from Streptomyces griseus (final concentration 0.1 U/mL; product # C9830, Sigma Aldrich) in a 1:1 (v/v) ratio for 6 h at 37 °C with gentle orbital shaking. The released EVs in the solution were filtered through Exo-spin columns (EX03 Exo-spin™ mini, Cell Guidance Systems) and analysed with a flow nanoanalyzer (NanoFCM) to determine particle concentration and percentage of EVs expressing transmembrane protein CD81 and/or cytosolic protein TSG101. The released vesicles were also imaged using cryogenic electron microscopy (cryo-EM). Uncoated EVs (∼1×10^9^ particles/mL) was used as a control.

The biological activity of released EVs was determined by measuring TNF-α production in macrophages. J774A.1 macrophages were seeded in 96-well plates (Thermo Fisher Scientific) at a density of 2×10^5^ cells/mL (0.1 mL/well) in culture medium alone and in culture medium containing 100 ng/mL lipopolysaccharide (LPS) from Escherichia coli O111:B4 (product #L2630, Sigma Aldrich) and 25 ng/mL interferon-𝛾 (IFN-𝛾) (product #GFH77, Cell Guidance Systems) to induce inflammation. The cells were incubated at 37 °C for 24 h. The next day, J774A.1 macrophages were treated with EVs released from the coating formulation (final concentration in wells: 10^9^ particles/mL) and incubated for a further 24 h. Uncoated MSC-EVs were used as a positive control for anti-inflammatory activity and cell culture medium was used as a negative control. TNF-α released from J774A.1 macrophages was quantified using a mouse Enzyme-Linked Immunosorbent Assay (ELISA; product #ELM-TNFa, RayBiotech), following the manufacturer’s protocol. The absorbance was measured at 450 nm using an Infinite 200 Pro plate reader (Tecan) and a standard curve was generated to quantify TNF-α levels in the samples.

### Effect of coated MSC-EVs in a DSS-induced colitis mouse model

All animal experiments were approved by the Institutional Animal Care and Use Committee (IACUC) of Nanyang Technological University (NTU). C57BL/6J mice (male, 8–10 weeks old, body weight ≥ 20 g) were obtained from the NTU animal research facility and housed under standard conditions with a 12-hour light/dark cycle and ad libitum access to food and water.

A total of 30 mice were randomly allocated to six groups (n = 5 per group). Mice in the healthy control group were housed together and given reverse osmosis (RO) water as standard drinking water for the duration of the study. Mice from the remaining five groups were co-housed to minimise cage effects. Acute colitis was induced by administering 3% (w/v) dextran sodium sulfate (DSS; 40 kDa MW, Sigma Aldrich, product #42867-100G, batch #BCCJ9094) in drinking water ad libitum for five consecutive days, followed by administration of normal drinking water. Clinical signs of colitis, including weight loss, diarrhoea, and occult blood in faeces, were recorded daily and scored using an adapted scoring system (Supplementary Figure 11). Scores for each parameter were summed to calculate the disease activity index (DAI). Treatments were administered once daily during DSS exposure (days 1–5). Mice received MSC-EVs (1×10^9^ particles/mouse/day) via intravenous (IV) injection in the tail vein or by oral gavage (100 µL/dose). For oral delivery, either uncoated or coated EVs in RO water were administered. Healthy and DSS control groups received RO water (n = 3) and saline injection (n = 2). An additional control group received coated liposomes in RO water orally. All preparations for *in vivo* administration were sterile-filtered or handled aseptically in a biosafety cabinet. Mice were euthanised by carbon dioxide inhalation and colons were excised and measured for length and weight in a blinded manner. Colons were also photographed on a flat surface and scale bars were generated in ImageJ from a ruler imaged in the same plane.

#### Biodistribution of MSC-EVs

MSC-EVs were labelled with Wheat Germ Agglutinin conjugated to Alexa Fluor 680 (WGA-AF680; #W32465, Thermo Fisher Scientific). Briefly, EVs were incubated with 0.1 mg/mL WGA-AF680 at 37 °C for 30 min under gentle agitation, protected from light. Labelled EVs were purified using Exo-spin columns (EX03 Exo-spin™ mini, Cell Guidance Systems). Mice received 3% DSS in drinking water for 4 days to induce colitis and EVs (1×10^10^ particles/mouse) were administer by IV injection or oral gavage (uncoated and coated EVs). After 24 h, fluorescence images of the colon, liver, spleen, heart, lungs, and kidneys were acquired using an *in vivo* imaging system (IVIS, Revvity) and analysed with *Living Image* software (v4.8.3) with background correction applied.

### Statistical analysis

All results are expressed as mean ± SD. GraphPad Prism^®^ 10 was used to construct graphs and perform statistical analysis. A p-value < 0.05 was considered statistically significant.

## Supplementary Data

Supplementary material is provided with this manuscript.

## Acknowledgements

We thank Giorgia Pastorin for her guidance and support throughout this work. The authors acknowledge the Advanced Flow Cytometry Platform, particularly Cynthia Bishop, and the Wohl Cellular Imaging Centre (WCIC) at King’s College London for instrument access and technical support. We also thank Matthias Wacker and his team for access to the NanoSight NS300 at the National University of Singapore (NUS); Lucas Lu for scanning electron microscopy data acquisition at NUS; Saw Wuan Geok for assistance with cryogenic electron microscopy at the NTU Institute of Structural Biology (NISB); Xavier Camous (Biomed Global) with imaging flow cytometry and Siobhan King (ONI) with super-resolution microscopy. We further acknowledge Evonik and Acoerela for providing Eudragit^®^ S-100 and liposomes, respectively. In vivo experiments were supported by a NAMIC Singapore grant (A-8001601-00-00).

## Author contributions

M.B. conceptualised the project, designed and conducted the experiments, and wrote and finalised the manuscript. W.H.C., R.P.K.M., Y.W.L., J.J., Y.Z. and X.L. performed and/or assisted with experiments. B.C. provided technical advice, resources and supervision. D.V. conceptualised the project, provided technical advice and resources, supervised the work, and edited the manuscript. All authors reviewed and approved the manuscript before submission.

## Data availability

The data that support the findings of this study are available from the corresponding author upon reasonable request.

## Supplementary material

**Supplementary Figure 1.**
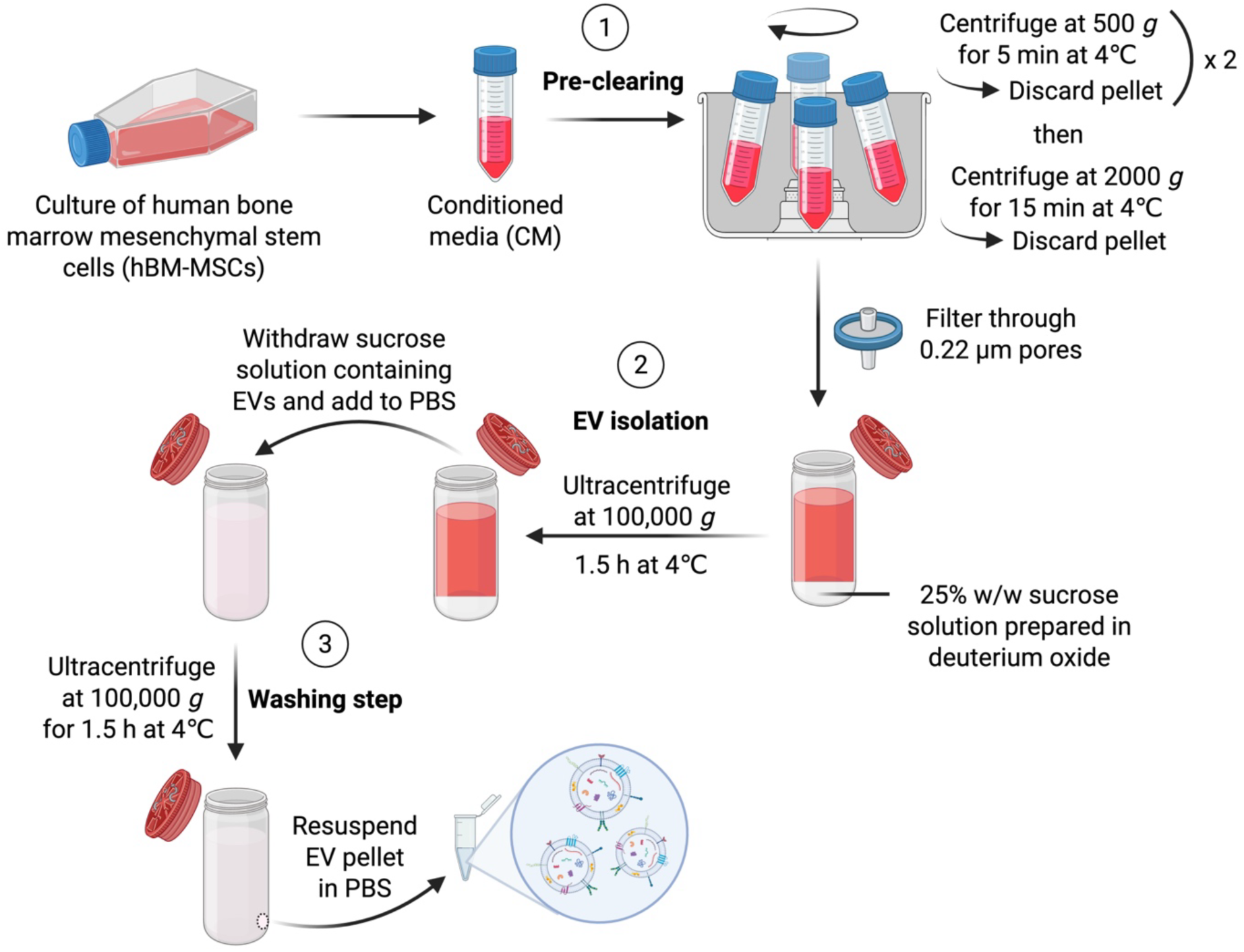
Isolation of EVs from human bone marrow mesenchymal stem cells. The sucrose cushion ultracentrifugation technique is a density-based isolation method that comprises three main steps: 1) Pre-clearing the conditioned medium (CM) from dead cells and cell debris. 2) Isolation of EVs from CM onto a sucrose cushion. 3) Washing step to remove sucrose and contaminating proteins. Created with BioRender.com.

**Supplementary Figure 2.**
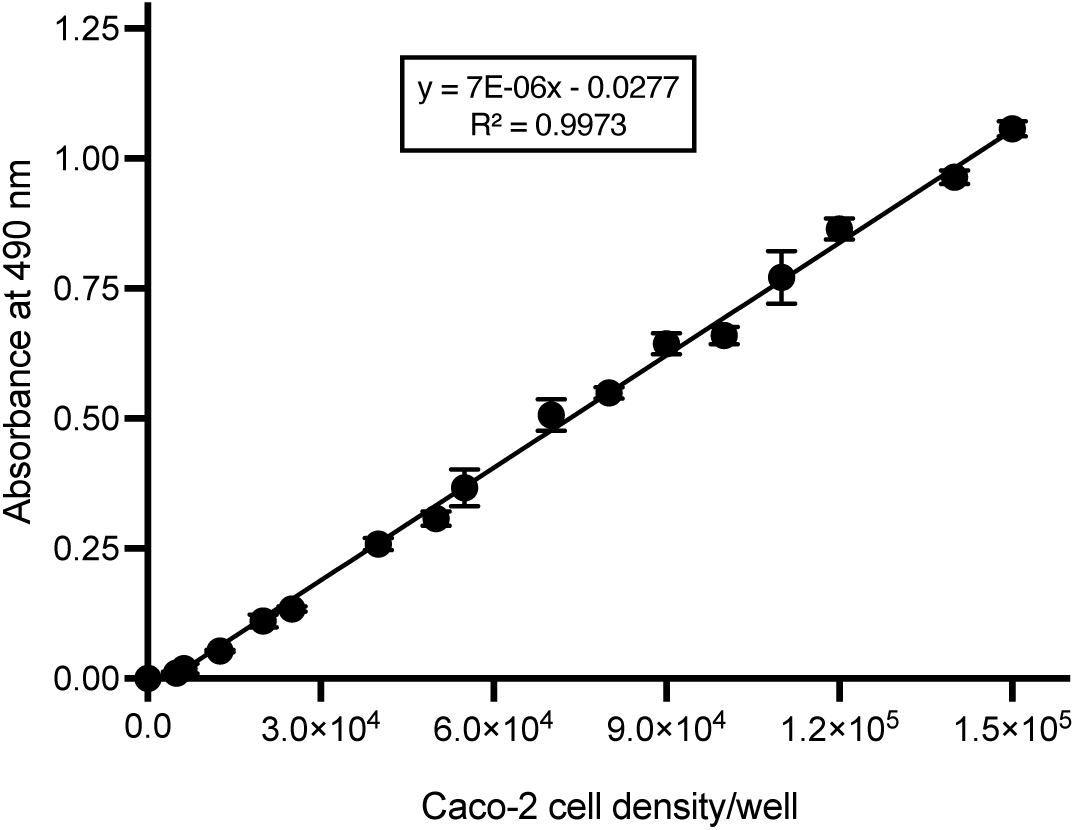
MTS assay standard curve. Caco-2 cells were seeded at different densities in 48-well plates and incubated overnight at 37 °C to allow cells to adhere to the wells. The next day, the wells were washed with PBS before 200 µL of serum-free medium and 20 µL of MTS reagent were added to the wells. After 2 h incubation at 37 °C, the absorbance of the wells at 490 nm was recorded using a plate reader (Infinite 200 Pro, Tecan). Absorbance of the formazan product at 490 nm is proportional to Caco-2 cell density/well. Each data point represents a mean of 3 replicates ± SD.

**Supplementary Figure 3.**
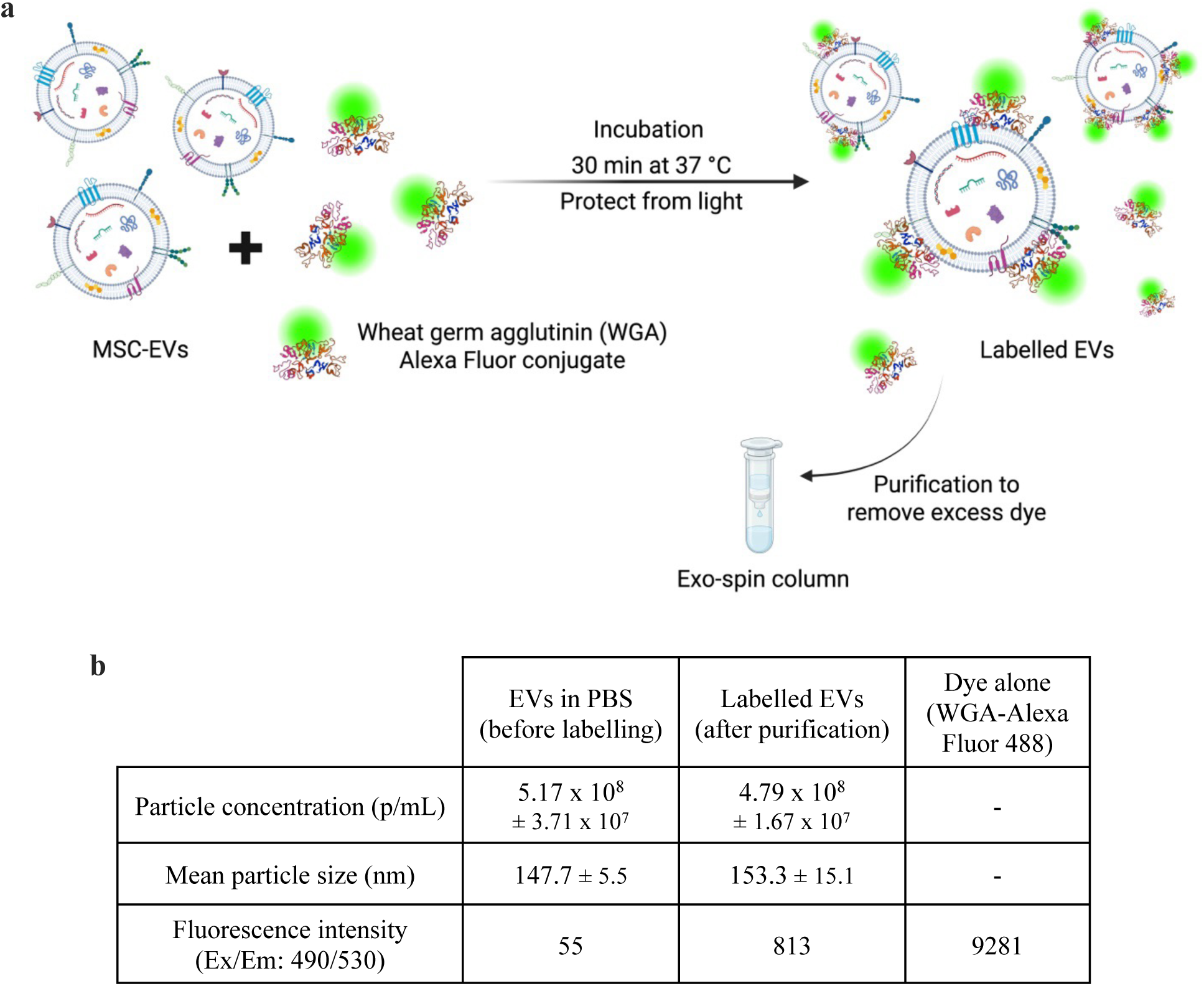
Labelling of MSC-EVs with Wheat Germ Agglutinin (WGA) Alexa Fluor conjugates. **(a)** MSC-EVs were incubated with WGA-Alexa Fluor 488 or 680 for 30 min at 37 °C with gentle agitation, protected from light. Excess dye was removed using Exo-spin^TM^ columns (Cell Guidance Systems) to purify the EVs. **(b)** Characterisation of MSC-EVs before and after fluorescent labelling. 90% of the EVs were recovered after labelling and purification, with no significant increase in the mean particle size. The size distribution and particle concentration were measured using nanoparticle tracking analysis (NTA) and the fluorescence intensity was measured using a plate reader (BioTek Synergy H1, Agilent).

**Supplementary Figure 4.**
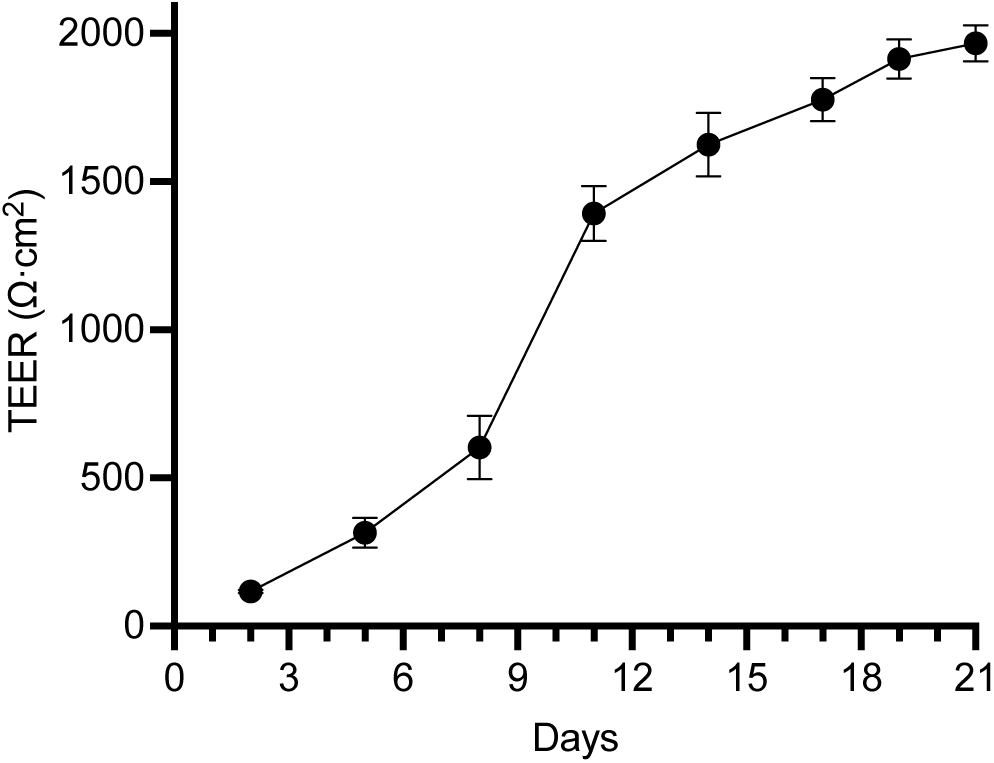
Transepithelial electrical resistance (TEER) measurements over 21 days. Colon Caco-2 epithelial cells were seeded on Transwell^®^ cell culture inserts and maintained in culture for 19-21 days until differentiation into intestinal epithelial cells and stabilisation of TEER values. TEER was measured every 2-3 days of the culture using an Epithelial Volt Ohm Meter (EVOM) to assess the development of barrier integrity and formation of tight junctions in Caco-2 monolayers. TEER values are expressed in ohm per cm^2^ (Ω·cm^2^). Each data point represents a mean of 5 independent replicates ± SD.

**Supplementary Figure 5.**
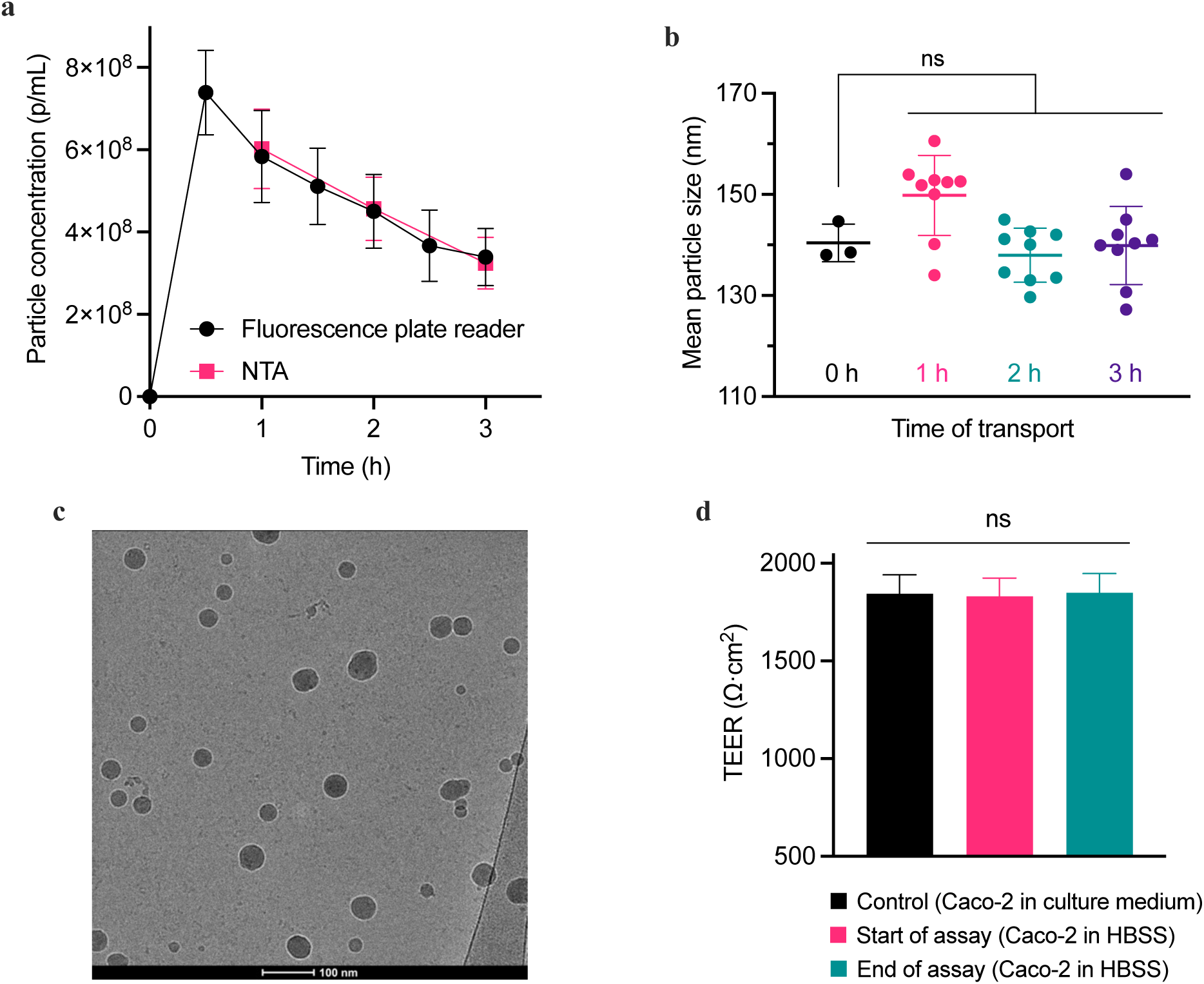
Transport of MSC-EVs across intestinal epithelial cells. Caco-2 cells were seeded on Transwell^®^ cell culture inserts and maintained for 19-21 days to allow differentiation into intestinal epithelial cells. The permeability assay was performed in HBSS. MSC-EVs labelled with WGA-Alexa Fluor 488 (1×10^9^ p/mL) were applied to the apical compartment of the Caco-2 monolayer and incubated at 37 °C for 3 h. **(a)** Particle concentration in the basolateral compartment of the Caco-2 monolayer over 3 h. Samples were collected every 30 min and measured using a fluorescence plate reader and nanoparticle tracking analysis (NTA). Most of the transport occurred during the first 30 min of the assay, followed by a decrease in the rate of transport as EVs reached an equilibrium across the semipermeable membrane. Internalisation of EVs by Caco-2 cells may also have contributed to reduced transport. **(b)** The mean particle size (measured by NTA) of MSC-EVs did not change significantly during transport, which suggests that the EVs remained intact after permeating the membrane (p > 0.05). **(c)** Cryogenic electron microscopy (cryo-EM) image of transported EVs showed structurally intact vesicles. Scale bar: 100 nm. **(d)** The transepithelial electrical resistance (TEER) values remained constant throughout the permeability study, indicating that the barrier integrity of Caco-2 cells was not affected by MSC-EVs (p > 0.05). Data are presented as mean ± SD (n = 3). **(b)** One-way ANOVA with post-hoc Dunnett’s test. **(d)** One-way ANOVA with post-hoc Tukey’s test. (* p > 0.05).

**Supplementary Figure 6.**
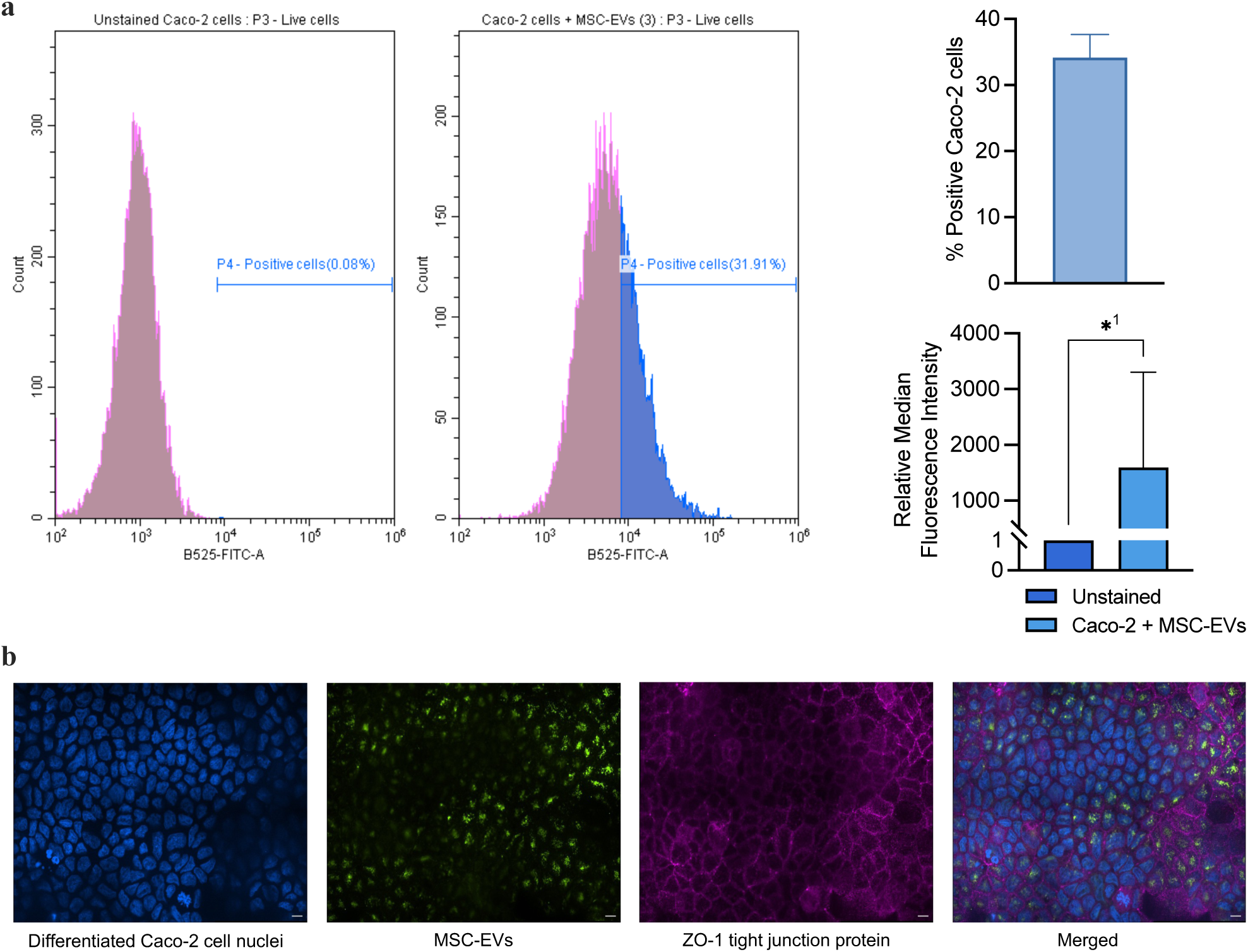
Uptake of MSC-EVs in differentiated Caco-2 intestinal epithelial cells. Differentiated Caco-2 cells were incubated with MSC-EVs labelled with WGA-Alexa Fluor 488. After 4 h, the cells were analysed using flow cytometry (Cytoflex LX, Beckman Coulter). **(a)** 35% of Caco-2 cells were positive for MSC-EVs and showed a significant increase in median fluorescence intensity (*^1^ p = 0.0453). Data are presented as mean ± SD (n = 2) with t-test (* p < 0.05). **(b)** Fluorescence images of MSC-EVs in intestinal epithelial cells. EVs were labelled with WGA-Alexa Fluor 488. Differentiated Caco-2 cell nuclei and tight junction protein ZO-1 were stained with DAPI and Alexa-Fluor 647, respectively. Images were captured using a Nikon Eclipse Ti Inverted Spinning Disk confocal microscope. Scale bar: 20 µm.

**Supplementary Figure 7.**
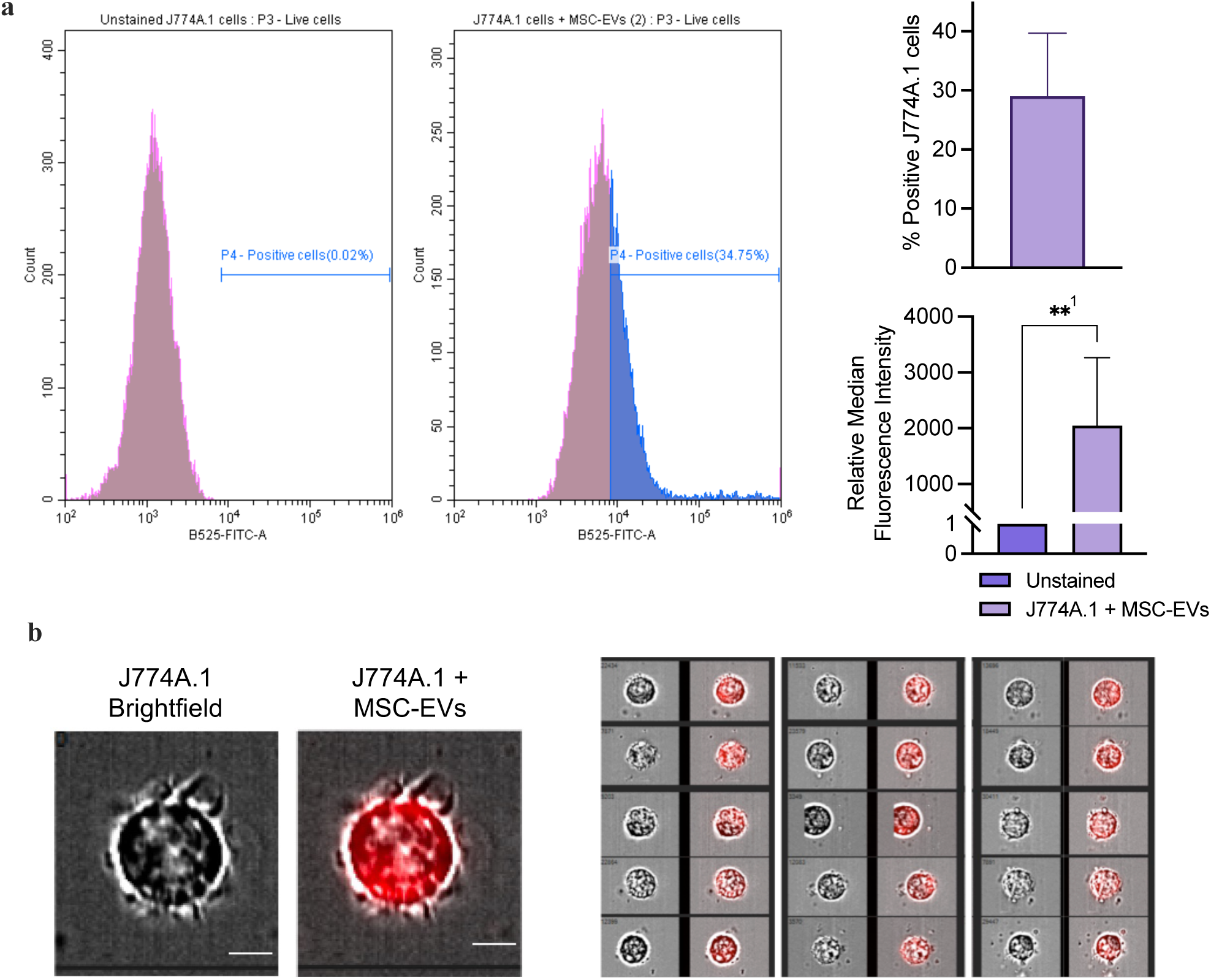
Uptake of MSC-EVs in J774A.1 macrophages. J774A.1 macrophages were incubated with MSC-EVs labelled with WGA-Alexa Fluor 488. After 4 h, the cells were analysed using flow cytometry (Cytoflex LX, Beckman Coulter). **(a)** 30% of J774A.1 cells were positive for MSC-EVs and showed a significant increase in median fluorescence intensity (**^1^ p = 0.0021). Data are presented as mean ± SD (n = 2) with t-test (** p < 0.01). **(b)** Fluorescence images of MSC-EVs in J774A.1 macrophages. EVs were labelled with WGA-Alexa Fluor 680. Images were captured using the Cytek Amnis ImageStream Mark II image flow cytometer. Scale bar: 5 µm.

**Supplementary Figure 8.**
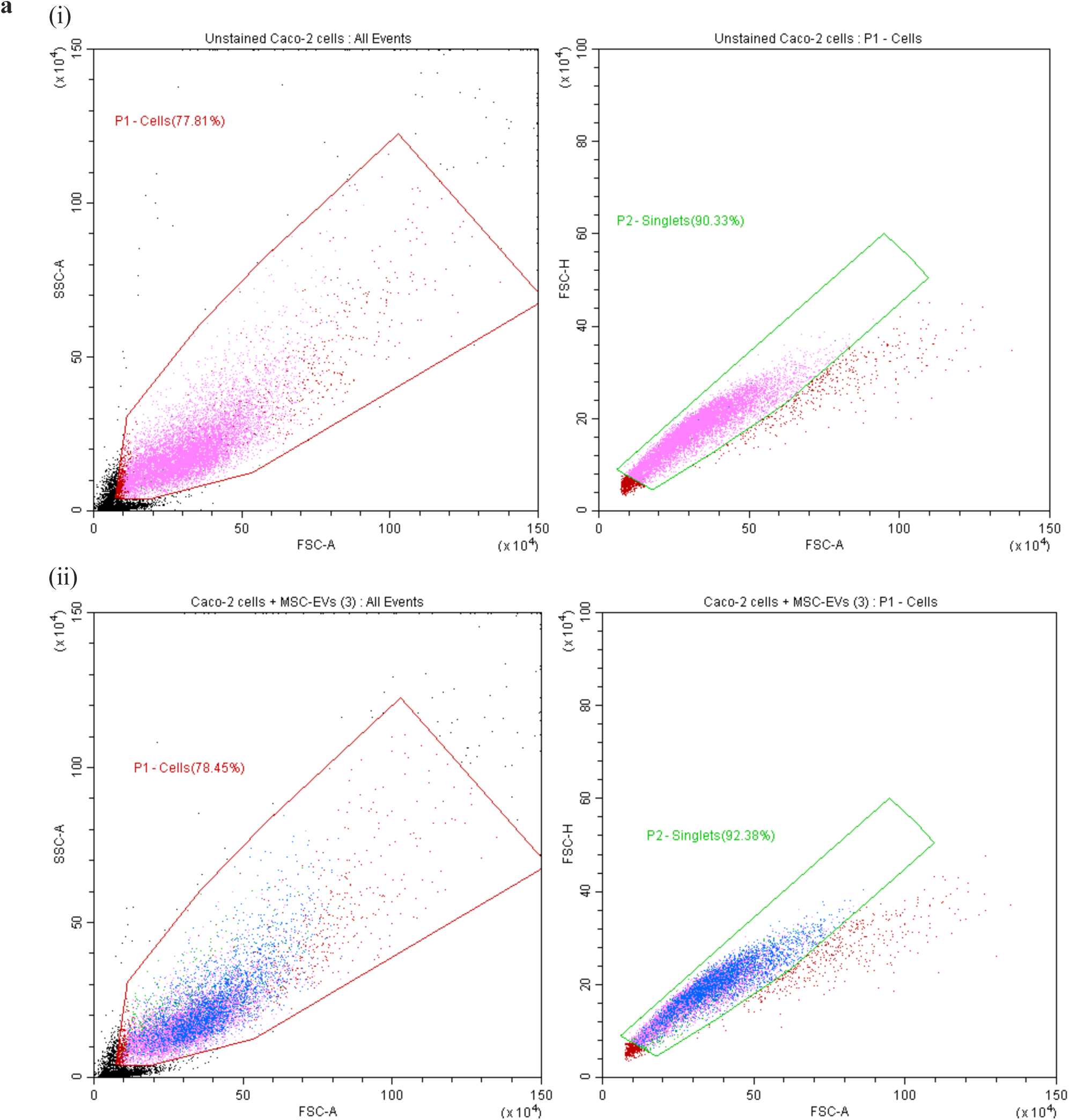
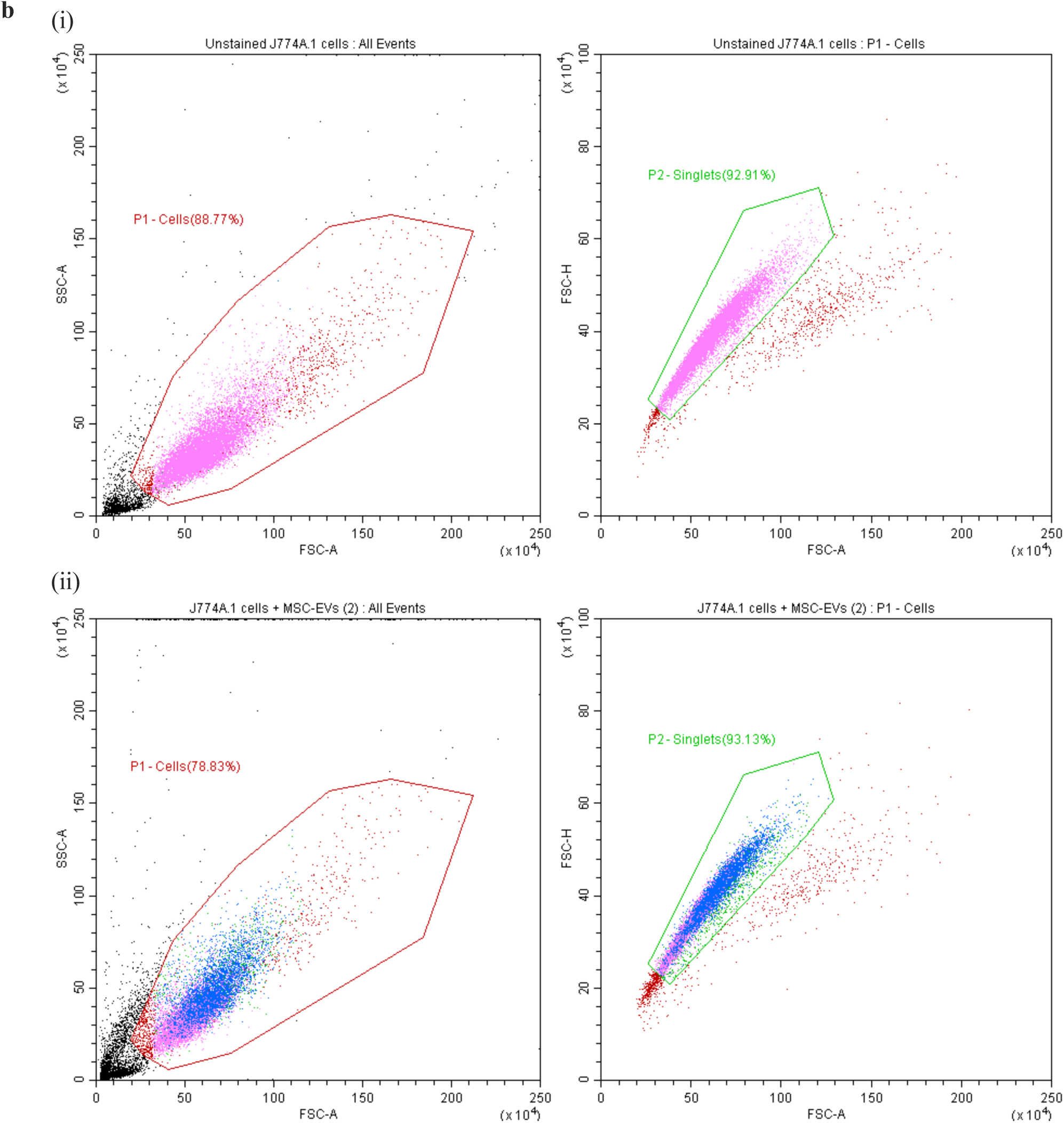
Gating used in flow cytometry analysis. Gating was manually set using the CytExpert software (Beckman Coulter) to measure the uptake of MSC-EVs in **(a)** differentiated Caco-2 intestinal epithelial cells, (i) Unstained cells, (ii) Positive cells and **(b)** J774A.1 macrophages, (i) Unstained cells, (ii) Positive cells.

**Supplementary Figure 9.**
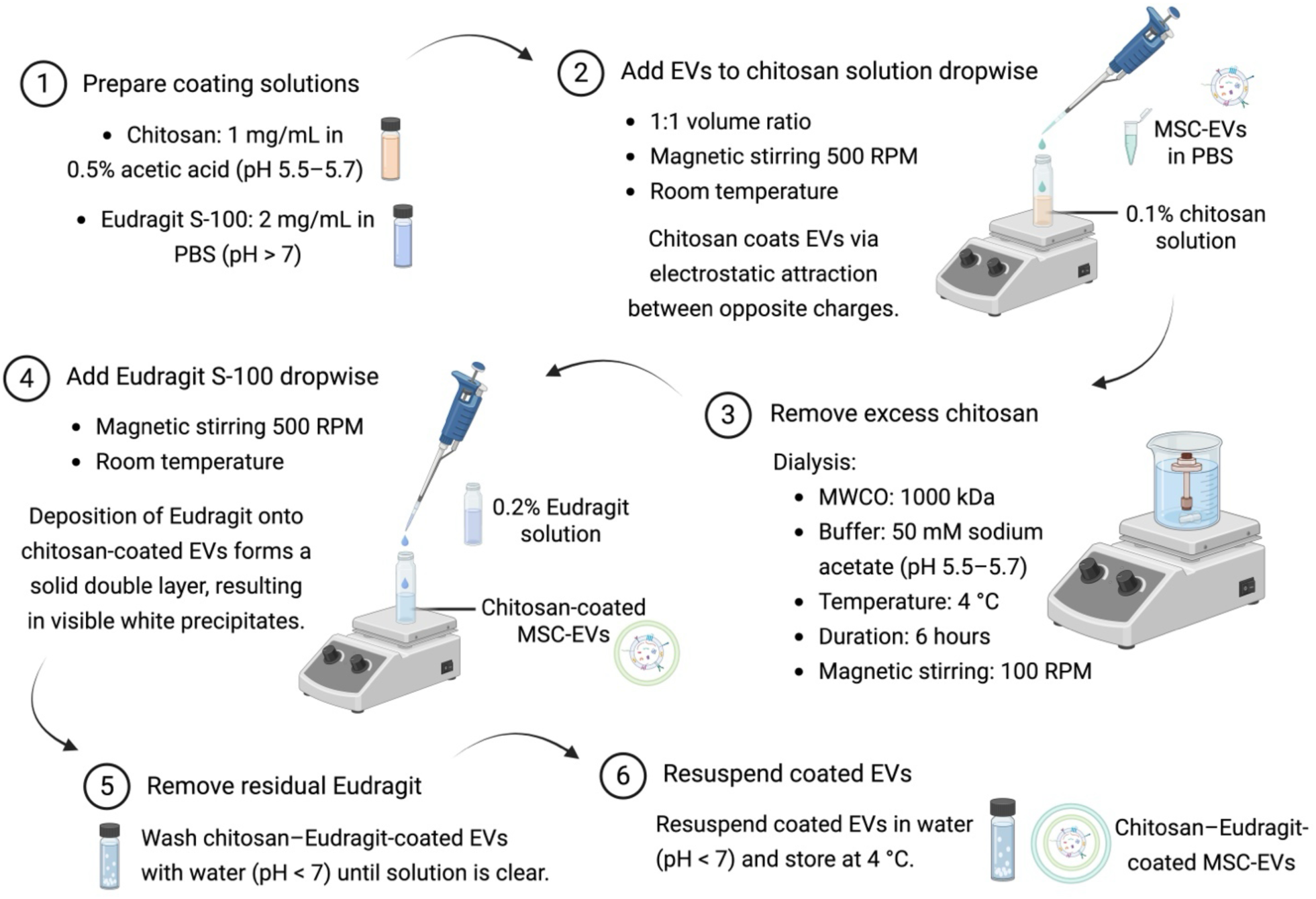
Preparation of coated MSC-EVs. MSC-EVs were sequentially coated with chitosan as the first layer and Eudragit S-100 as the second layer, resulting in chitosan–Eudragit-coated EVs (referred to as coated EVs). Created with BioRender.com.

**Supplementary Figure 10.**
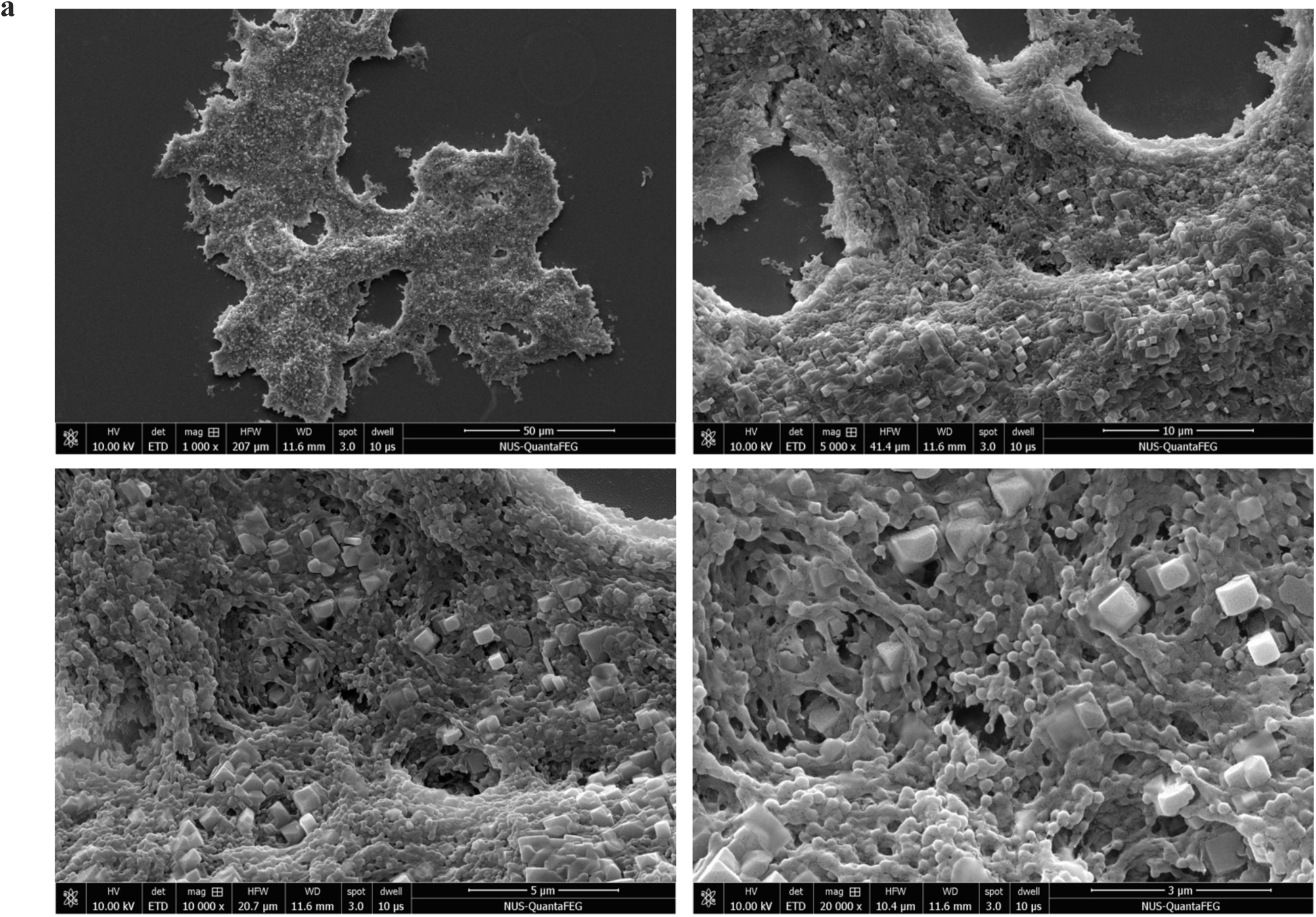
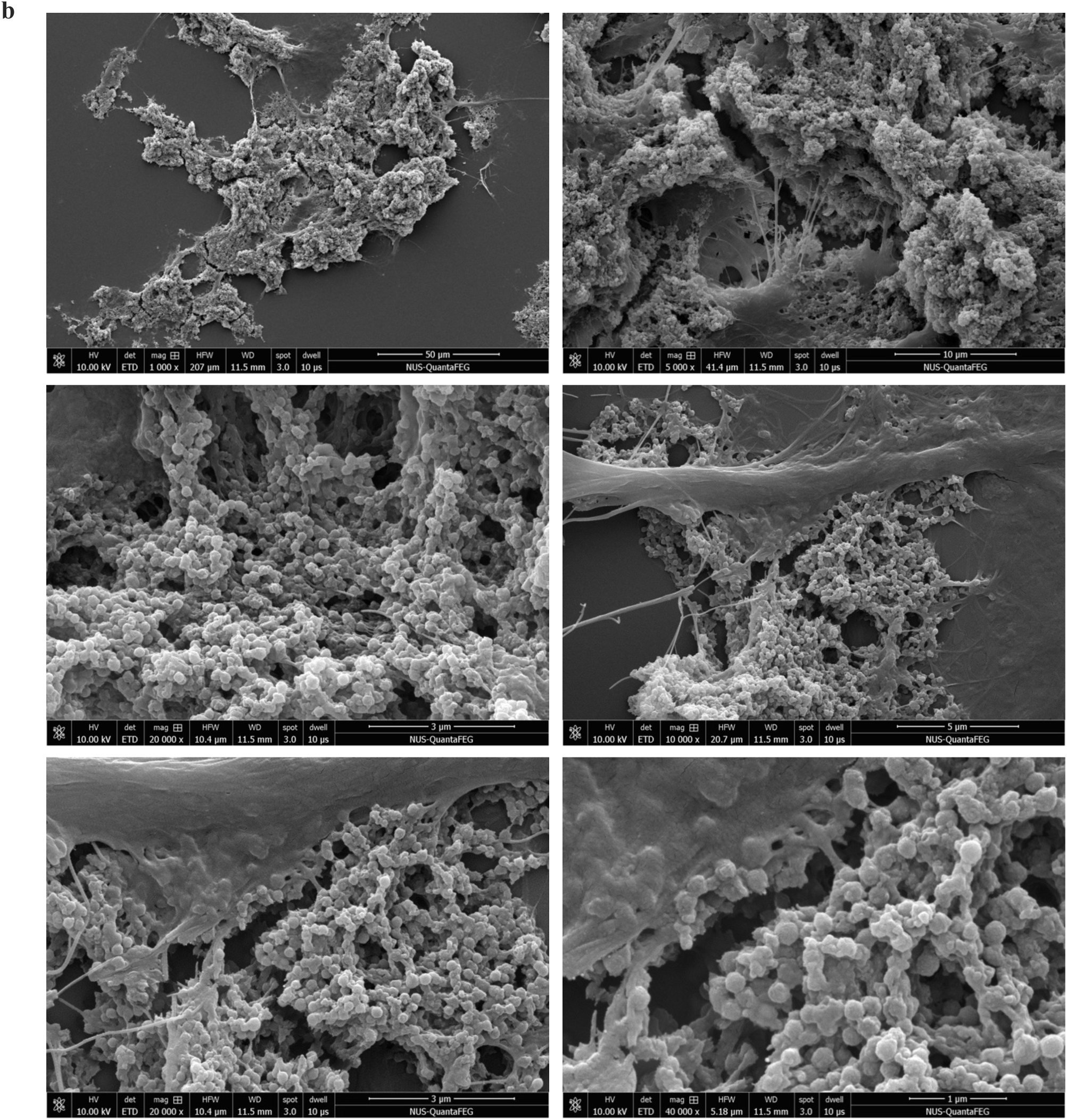
Scanning electron microscopy (SEM) images of coated MSC-EVs. **(a)** in water (pH 6.5) and **(b)** after incubation with gastric and intestinal simulated fluids and digestive enzymes (pepsin and pancreatin).

**Supplementary Figure 11.**
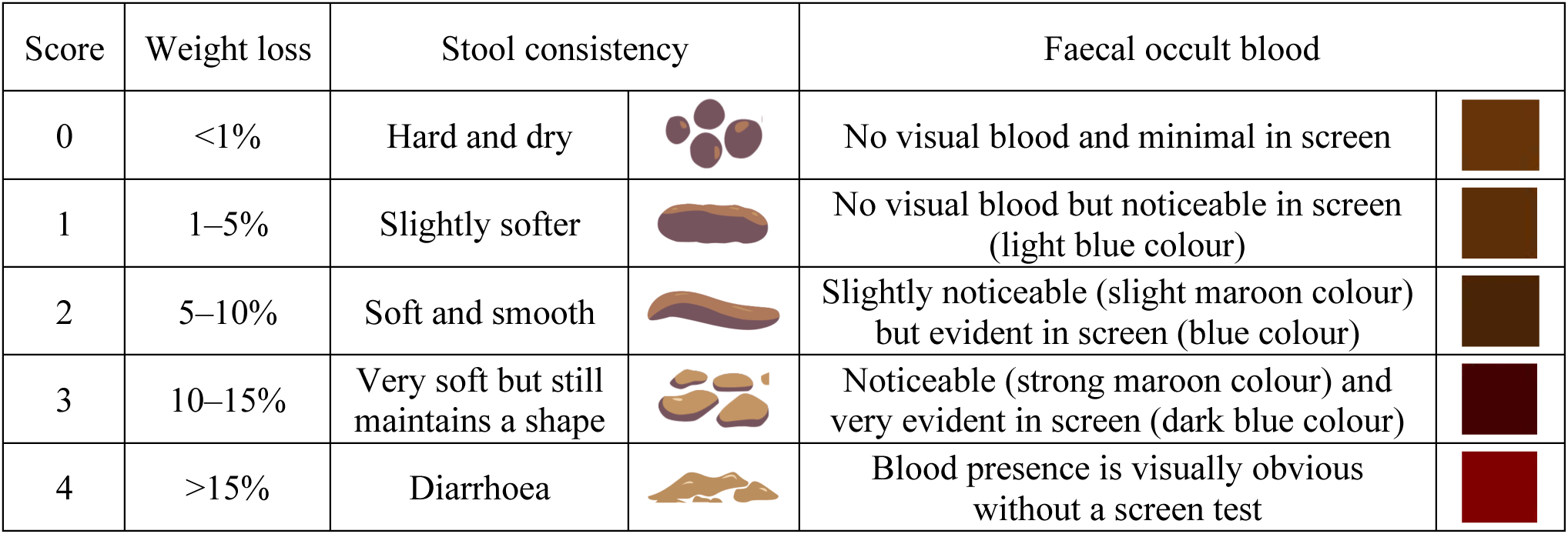
Adapted scoring guide for body weight loss, stool consistency and faecal occult blood. Blood presence in stools was assessed using Hema-Screen^®^ test.

**Supplementary Figure 12.**
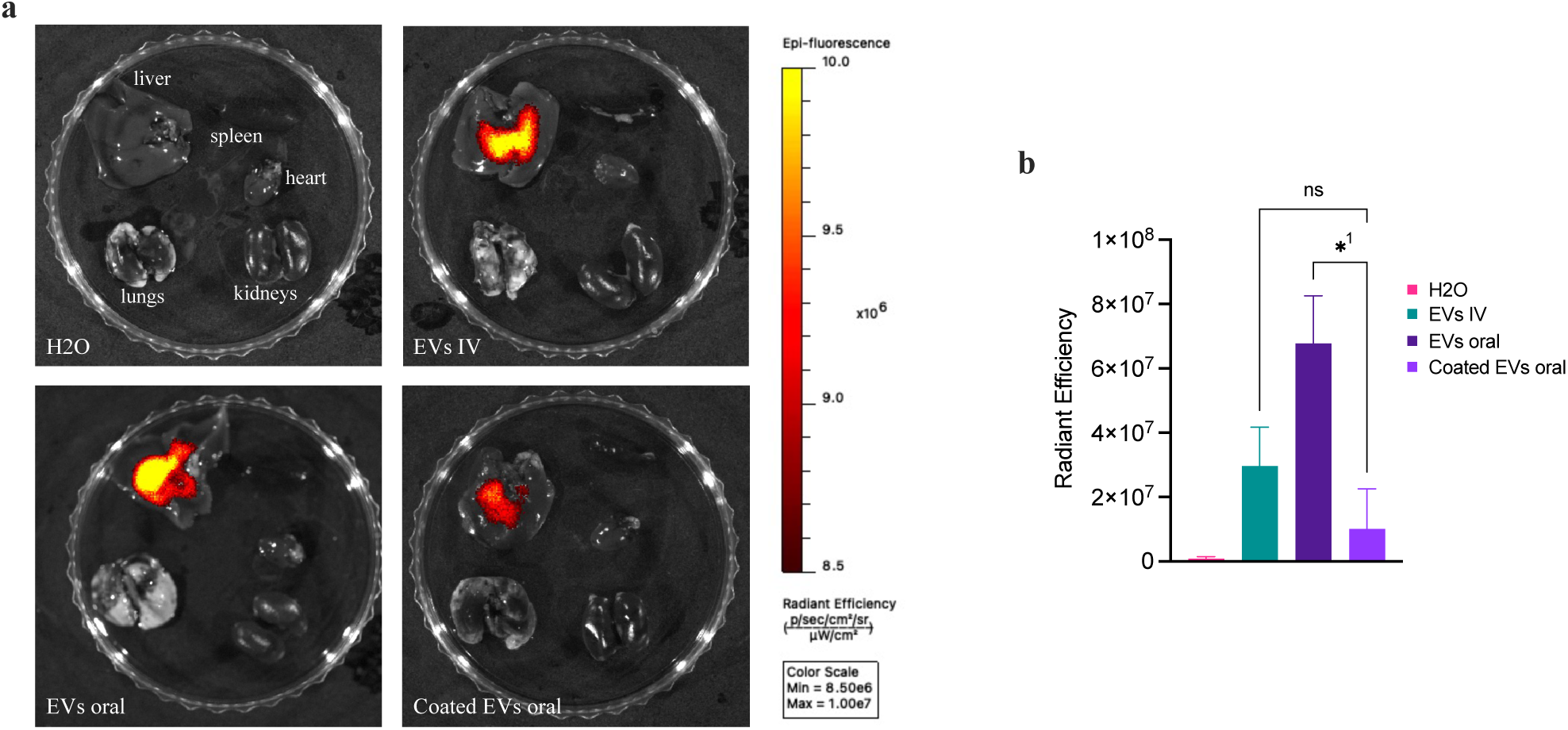
Biodistribution of MSC-EVs in the liver. **(a)** Biodistribution of MSC-EVs in the liver, spleen, heart, lungs and kidneys of DSS mice 24 h post dosing. WGA-AF680–labelled EVs were administered via IV injection and oral gavage for uncoated and coated EVs and the fluorescence in the organs was imaged using IVIS (Revvity). **(b)** Quantification of EV fluorescence signal (radiant efficiency) in the liver 24 h after administration. Uncoated EVs administered orally (EVs oral) showed significantly higher fluorescence intensity in the liver compared with coated EVs (^1^* p = 0.0166). Data are presented as mean ± SD (n = 2) with one-way ANOVA (* p < 0.05).

